# Protective host-dependent antagonism among *Pseudomonas* in the *Arabidopsis* phyllosphere

**DOI:** 10.1101/2021.04.08.438928

**Authors:** Or Shalev, Talia L. Karasov, Derek S. Lundberg, Haim Ashkenazy, Detlef Weigel

## Abstract

The plant microbiome is a rich biotic environment, comprising numerous taxa. The community structure of these colonizers is constrained by multiple factors, including host-microbe and microbe-microbe interactions, as well as the interplay between the two. While much can be learned from pairwise relationships between individual hosts and microbes, or individual microbes with themselves, the ensemble of interrelations between the host and microbial consortia may lead to different outcomes that are not easily predicted from the individual interactions. Their study can thus provide new insights into the complex relationship between plants and microbes. Of particular importance is how strain-specific such plant-microbe-microbe interactions are, and how they eventually affect plant health. Here, we test strain-level interactions in the phyllosphere between groups of co-existing commensal and pathogenic *Pseudomonas* among each other and with *A. thaliana*, by employing synthetic communities of genome-barcoded isolates. We found that commensal *Pseudomonas* prompted a host response leading to a selective inhibition of a specific pathogenic lineage, resulting in plant protection. The extent of plant protection, however, was dependent on plant genotype, indicating that these effects are host-mediated. There were similar genotype-specific effects on the microbe side, as we could pinpoint an individual *Pseudomonas* isolate as the predominant cause for this differential interaction. Collectively, our work highlights how within-species genetic differences on both the host and microbe side can have profound effects on host-microbe-microbe dynamics. The paradigm that we have established provides a platform for the study of host-dependent microbe-microbe competition and cooperation in the *A. thaliana*-*Pseudomonas* system.

## Introduction

Plants, like other complex organisms, host a diverse set of microbes. The assembly of these microbial communities is shaped both by host-microbe as well as microbe-microbe interactions. These interactions may be of any symbiotic type: mutualistic, commensalistic or parasitic, and are dictated by the balance of inhibition and facilitation of growth. As has been exemplified in many studies, interactions between organisms are not static, but rather a dynamic process that depends on the environment - both biotic [1,2] and abiotic [3,4] - as well as on evolutionary history [5,6].

Many aspects of the dynamic interactions between plants and microbes have been studied in considerable detail, not least because of their implications for agriculture and ecology. Colonization of the plant depends on the ability of microbes to grow on and in the host, but also on the antagonistic ability of the host to promote or restrict such microbial growth. In the case of pathogens, there is often a co-evolutionary arms race, in which plants evolve recognition and immune tools to restrict the growth of microbes, while microbes evolve evasion and an offensive arsenal to further populate the plant [7,8]. These co-evolutionary dynamics typically fuel the generation of genetic diversity within both host and microbe, and the dependence of microbial colonization and host health on intraspecific variation has been documented in numerous studies [5,9–11]. Nonetheless, the extent to which intraspecific host variation shapes overall microbial composition is minimal [3,4], with the most dramatic effects seen for specific taxa that are recognized by the immune system [12,13]. Instead, other environmental factors have a much larger influence on the overall composition [3,4], including other resident microbes [1,14,15]. Taken together, this suggests that successful colonizers reflect compatibility to grow in the presence of both the host and other microbes, and that this compatibility depends on their genetic makeup.

The colonizing microbes exert differential effects on host health - from harmful [16] to beneficial [17]. These effects are mainly related to microbial load, since overpopulation of the plant by microbes can negatively impact its health [9,18]. Nonetheless, the host has the genetic arsenal to control the growth of some microbes, thus avoiding negative outcomes [7]. This raises questions about the ability of the host plant to differentially recognize and respond to a consortium of microbes with a range of functions, i.e. differentiating friend from foe in a complex assembly of microbial taxa. While there is a growing body of literature about host response to individual pathogens [19] and individual commensals [20], a more realistic scenario is the integrated host response to communities that include both diverse pathogens and diverse commensals.

In the same way, the numerous constraints resulting from multiple host-microbe and microbe-microbe interrelations create a complex system of relationships, making extrapolation of rules from simplistic systems likely difficult. For example, overpopulation of the plant by one microbe can result in negative health impacts, but these might be mitigated in the presence of other microbes [17,21]. While studies of microbe-microbe interactions *in planta* have paved the way for important findings about their impact on the overall community [14,15], the effect of the host on such microbial interactions has often not been considered, despite the host being able to affect these via direct host-microbe interactions [22]. Hence, the high degree of interconnectedness at the host-microbe-microbe interface calls for holistic research of this system, rather than tackling individual components, to unravel dynamics that result from the multiple constraints. Such an approach can be conducted using synthetic communities, which establish causality and not only associations between microbe-microbe and plant-microbe interactions [23].

In a previous study, Karasov and colleagues [11] surveyed *Pseudomonas* populations from leaves of wild *Arabidopsis thaliana* plants in south-west Germany. Among these, one lineage, which was highly pathogenic in axenic infections, often dominated endophytic microbial communities of *A. thaliana* leaves. Nonetheless, this lineage was isolated from plants without any visible disease symptoms, suggesting that other factors, including co-colonizing microbes, were mitigating the pathogenic phenotype. This includes other *Pseudomonas* lineages, which did not appear to have any significant impacts on host health when tested individually [11].

Here, we took advantage of our collection of wild *Pseudomonas* isolates to investigate intraspecific host-microbe-microbe dynamics by infecting *A. thaliana* plants with synthetic *Pseudomonas* communities. Specifically, we examined interactions between pathogenic and commensal *Pseudomonas* with the host leaves and with themselves, and the linkage of these to the host health. We found that the host facilitated protective commensal-pathogen interactions, and revealed further complex interactions that could not be realized by studying host-microbe or microbe-microbe relationships individually.

## Results

### Genome barcoding of *Pseudomonas* isolates and experimental design

To test possible host-commensal-pathogen dynamics in local populations, we colonized six *A. thaliana* genotypes with synthetic bacterial communities composed of pathogenic and commensal *Pseudomonas* candidates. Pathogenicity classification was based on demonstrated pathogenic potential effects of different *Pseudomonas* lineages in the Karasov collection [11]. Only one lineage - which dominated local plant population - was associated with pathogenicity, both according to its negative impact on rosette weight and to visible disease symptoms [11]. This lineage was previously named “OTU5” (Operational Taxonomic Unit number 5) [11]. We henceforth call “ATUE5” (isolates sampled from Around TUEbingen, group 5) to all isolates that share a common 16S rDNA sequence in the V3-V4 region, previously associated with pathogenicity, and “non-ATUE5” to all other *Pseudomonas* from the Karasov collection [11]. We interchangeably use the terms pathogens and ATUE5, as well as commensals and non-ATUE5. We used host genotypes that originated from the same host populations from which the Pseudomonads were isolated - neary Tübingen, Germany (**Figure 1A**), aiming to reflect interactions between coexisting hosts and microbes.

**Figure 1.**
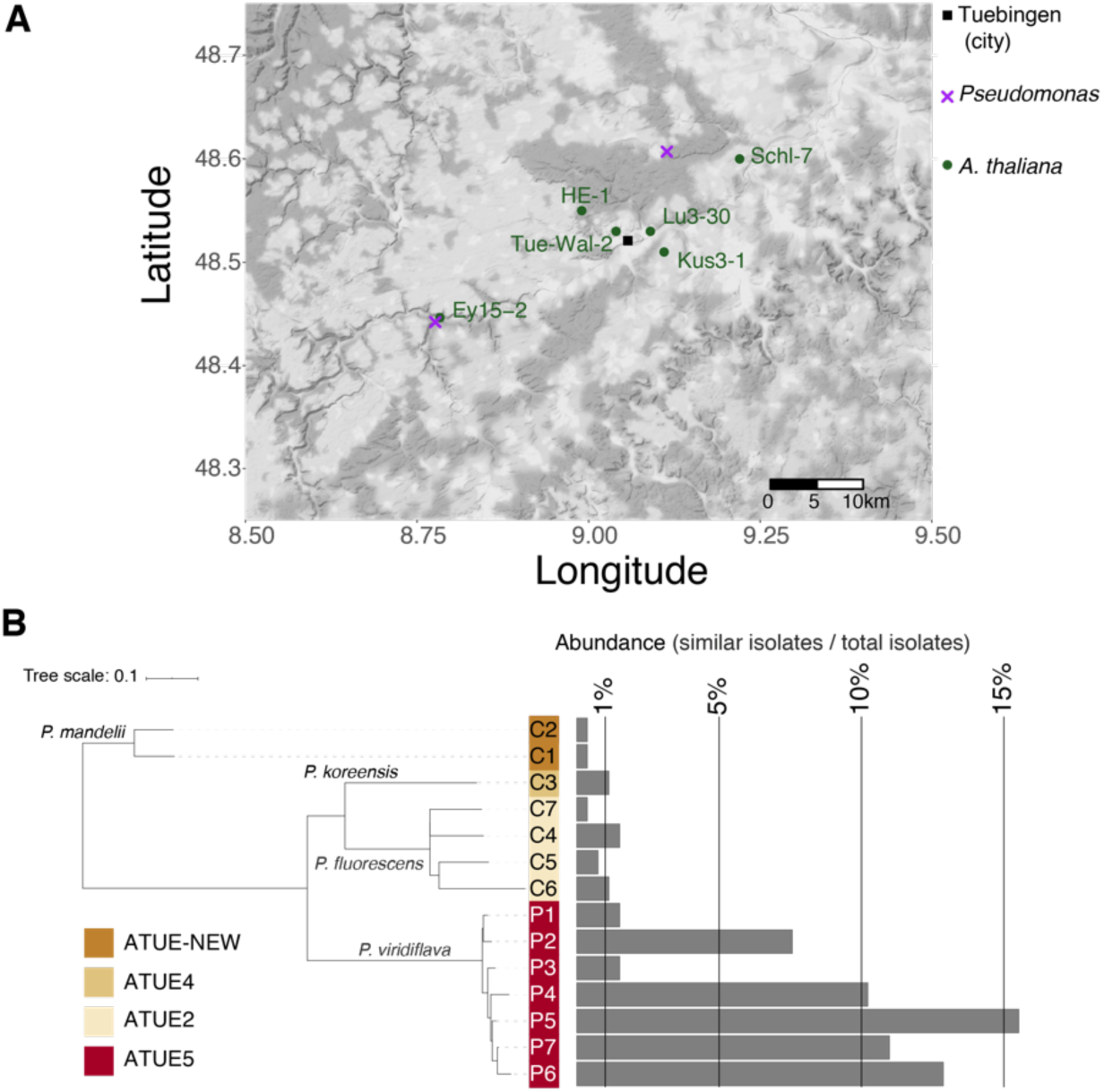
Study system. **A.** Location of original *A. thaliana* and *Pseudomonas* sampling sites around Tübingen (Germany). **B.** Taxonomic representation of the 14 *Pseudomonas* isolates used, and their respective abundance in the 1,524 strains of the Karasov collection [11]. Isolates were binned according to similarity (divergence < 0.0001 in core genome). Taxonomic assignment is indicated for each ATUE group (corresponding to a specific OTU in [11]). ‘P’ - Pathogen candidate. ‘C’ - Commensal candidate.

Overall, seven pathogenic *Pseudomonas* and seven commensal isolates were chosen, prioritizing those with the highest estimated abundance in the field (**Figure 1B**). The abundance was estimated by the number of similar isolates (defined as nucleotide sequence divergence less than 0.0001 in their core genome) sampled in the original survey. Thus, the chosen isolates act as representatives for other similar isolates. In total, all 14 *Pseudomonas* isolates were classified as belonging to four OTUs, following 16S rDNA clustering at 99% sequence identity. Because of the high relatedness of several of the isolates, we could not rely upon a single endogenous genetic marker to distinguish between isolates. Instead, we genome-barcoded each of the isolates. We employed the mini-Tn7 system [24] to insert a single-copy of a 22 bp long unique sequence, flanked by universal priming sites, into the chromosome of each isolate (Illustration in **Figure S1A**). We validated the sequence of all barcodes in the corresponding isolates using Sanger sequencing (**Table S1**), and confirmed barcode integration by barcode-specific PCR (**Figure S2A**). Furthermore, we confirmed that barcode-amplification yielded the expected products when PCR-amplified from DNA extracted from infected *A. thaliana* individuals (**Figure S2B**). While barcoding slightly impaired the growth rates of the isolates P3 and P4, the majority of barcoded bacteria exhibited similar growth dynamics as the non-barcoded parental strains when tested in Lysogeny Broth (LB) medium (**Figure S3**).

Next, we constructed three synthetic communities using the barcoded isolates: An exclusively pathogenic synthetic community, comprising the seven ATUE5 isolates (hereafter ‘PathoCom’), an exclusively commensal synthetic community, comprising the seven non-ATUE5 isolates (hereafter ‘CommenCom’), and a joint synthetic community comprising all 14 isolates - both pathogens and commensals (hereafter ‘MixedCom’). Isolates were mixed in an equimolar fashion, and their absolute starting concentration was identical in each synthetic community. Thus, the inoculum of the MixedCom with 14 isolates had twice the total number of bacterial cells as either the PathoCom or CommenCom inoculum.

The community experiments were conducted in plants grown on soil in the presence of other microbes. Our decision to perform experiments on non-sterile soil stemmed from initial observations that the infection outcomes of plants grown on soil were more consistent with the outcomes observed in the field than infections of axenically grown plants. Specifically, our initial isolation of the focal bacterial strains was done from plants in the field that were alive and not obviously diseased [11]. In the lab, axenic infections with these strains showed rapid and dramatic phenotypic effects on the plants, often killing the plants as early as three days-post-infection (**Figure S4**). In contrast, soil-grown plants displayed only mild disease symptoms and decreased size 12 dpi (**Figure S4**), phenotypes more consistent with those observed in the field.

To more closely mimic natural infections, which likely occur through the air, we chose to infect plants by spraying with bacterial suspension, rather than direct leaf infiltration, as is common for testing of leaf pathogenic bacteria in *A. thaliana*. Twenty one days after sowing, we spray-infected the leaves of soil-grown *A. thaliana* plants raised in growth chambers with the three synthetic communities, and with bacteria-free buffer (hereafter ‘Control’). Twelve days after infection (dpi), we sampled the fresh rosettes, weighed them and extracted DNA (see Methods). Subsequently, we coupled barcode-specific PCR and qPCR. We included an amplicon from an *A. thaliana*-specific genomic sequence in the qPCR assay, which allowed us to approximate the absolute abundance per isolate, i.e., the ratio of isolate genome copies to plant genome copies (**Figure S1B**).

### Host-genotype effects on composition of synthetic communities

The six *A. thaliana* genotypes used in this study were originally sampled from the same geographic region (**Figure 1A**) - a maximum of 40 km apart. They were all from the area from which the *Pseudomonas* strains used were isolated [11], and they were also all from the same host genetic group [25]. In accordance, we expected that host genotype would have little, if any effect on the composition of our synthetic communities of local *Pseudomonas* isolates. However, while not large, there was a significant effect of host genotype, explaining 5 to 12% of compositional variation in the different communities, as determined by permutational multivariate analysis of variance (PERMANOVA) with Bray–Curtis distances (**Table 1**). For comparison, the batch effect (between the different experiments) explained up to 26% of compositional variation.

**Table 1.** Permutational multivariate analysis of variance (PERMANOVA) based on Bray-Curtis distances, for compositions of the 14 barcoded bacteria in treated hosts. The analysis was constrained by the host genotype and the experiment batch (‘exp’) to estimate their effect on the explained variance.

| Treatment | Df | Sum Sq | Pseudo-F | R2 | Pr(>f) | Variation source |
| --- | --- | --- | --- | --- | --- | --- |
| PathoCom | 5 | 4.83 | 4.67 | 0.1199 | **0.0005** | Genotype |
|  | 1 | 1.68 | 8.11 | 0.0417 | **0.0005** | Exp |
|  | 5 | 1.08 | 1.04 | 0.0268 | 0.3973 | Genotype:Exp |
|  | 158 | 32.68 | NA | 0.8116 | NA | Residuals |
|  | 169 | 40.27 | NA | 1.0000 | NA | Total |
| CommenCom | 5 | 2.36 | 3.88 | 0.0839 | **0.0005** | Genotype |
|  | 1 | 7.32 | 60.19 | 0.2604 | **0.0005** | Exp |
|  | 5 | 1.53 | 2.52 | 0.0545 | **0.0030** | Genotype:Exp |
|  | 139 | 16.89 | NA | 0.6012 | NA | Residuals |
|  | 150 | 28.1 | NA | 1.0000 | NA | Total |
| MixedCom | 5 | 2.17 | 1.99 | 0.0456 | **0.0020** | Genotype |
|  | 1 | 6.29 | 28.89 | 0.1324 | **0.0005** | Exp |
|  | 5 | 2.03 | 1.86 | 0.0427 | **0.0020** | Genotype:Exp |
|  | 170 | 36.99 | NA | 0.7793 | NA | Residuals |
|  | 181 | 47.47 | NA | 1.0000 | NA | Total |

In bold, statistically significant relationships ( $P \leq 0.05$ ).

Analysis of similarities (ANOSIM) within each experiment indicated similar trends as PERMANOVA - with the genotype having a significant effect on isolate composition in each synthetic community (**Table S2A**).

We then examined bacterial composition clustering according to host genotype, by applying multilevel pairwise comparison using adonis (pairwise adonis, based on Bray-Curtis distances). Some pairs of genotypes differed in their effects on all three communities (**Table S2B**), an observation that was supported by nonmetric multidimensional scaling (NMDS) ordination of bacterial composition in each treatment (**Figure S5A**). The cumulative load of all isolates was associated with the loading on the NMDS1 axis (Pearson’s r > 0.99 and p-value < 2.2e^−16^, for all three communities), suggesting that a part of the compositional differences between host genotypes was due to absolute rather than relative abundance. In agreement, we observed differences in total bacterial load between the host genotypes, and the nature of the differences was treatment-dependent (**Figure S5B**).

How do these different community compositions affect plant growth?

### Host-genotype dependent pathogenicity, growth promotion or protection

PathoCom infection caused plants to grow less than control plants, during the 12 days of the experiment (**Figure 2**; **Figure S6**). In two out of the six host genotypes - Lu3-30 and TueWal-2 - weight decrease was milder, indicating a certain level of resistance to the PathoCom members (mean difference to control: Lu3-30 −29.1 mg [−59.3, −1.4], TueWal-2 −30.0 mg [−46.4, −13.4], Kus3-1-77.2 mg [96.4, 54.2], Schl-7 −93.1 mg [123.5, 67.7], Ey15-2 −92.5 mg [116.4, 66.0] and HE-1 - 53.9 mg [82.6, 27.0], with 95% confidence intervals in brackets). To validate that the effect of the PathoCom on plant weight was due to bacterial activity, and not merely a host response to the inoculum (e.g., PAMP-triggered immunity), we infected plants with heat-killed PathoCom. We found a minor weight decrease in three out of the six genotypes, but the overall contribution to weight reduction was small (**Figure S7**; heat-killed PathoCom accounts for 14% of the variation explained by the living PathoCom in the model shown).

**Figure 2.**
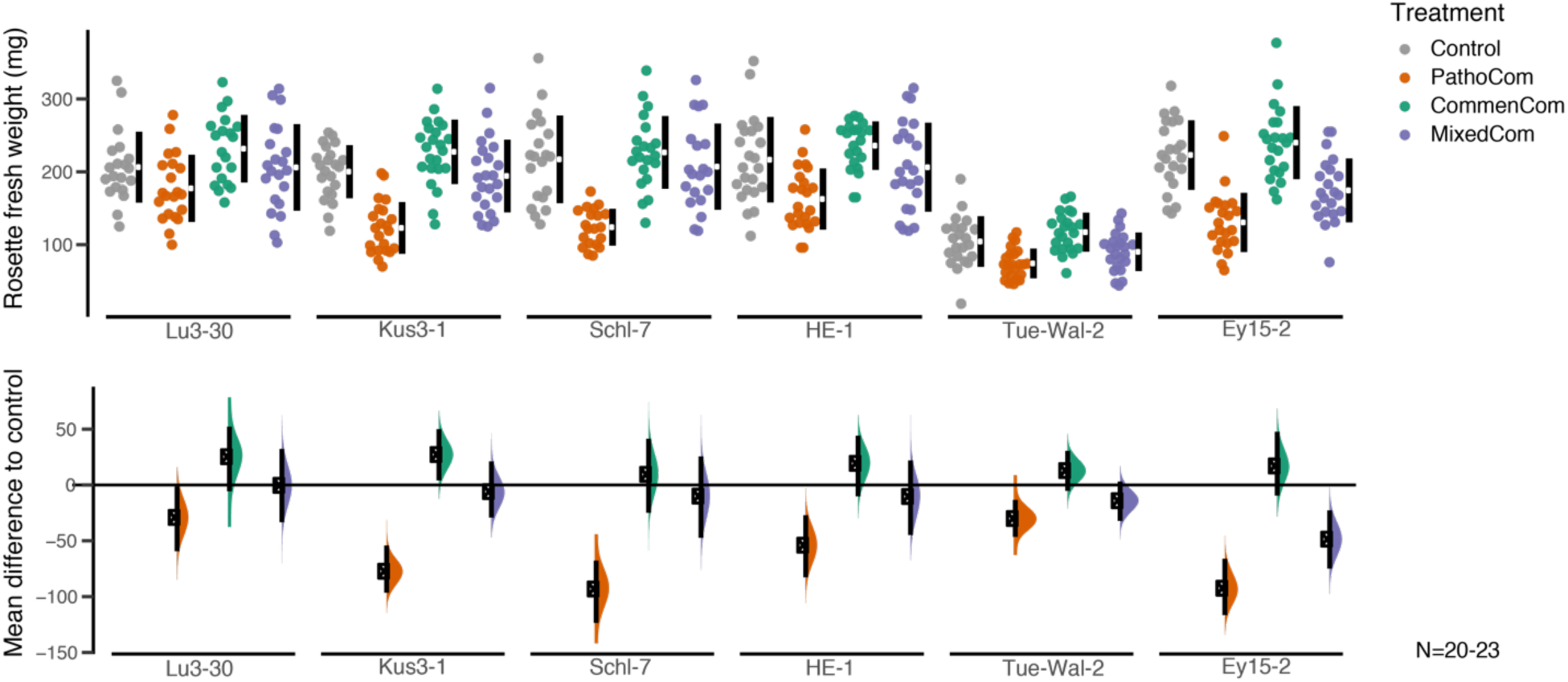
Commensal *Pseudomonas* protect the plant in a host-dependent manner. Each of the six *A. thaliana* genotypes used in this study was treated with Control, PathoCom, CommenCom and MixedCom. Fresh rosette weight was measured 12 dpi. The top panel presents the raw data, the breaks in the black vertical lines denote the mean value of each group, and the vertical lines themselves indicate standard deviation. The lower panel presents the mean difference to control, inferred from bootstrap sampling [26][27], indicating the distribution of effect sizes that are compatible with the data. 95% confidence intervals are indicated by the black vertical bars. Shown here are the results of one experiment. A second experiment gave similar results.

In contrast to PathoCom, infections with CommenCom led to a slight increase in fresh weight, suggesting plant growth promotion activity or alternatively protection from resident environmental pathogens (**Figure S6A**). This effect was independent on the host genotype (**Figure S6B**).

Importantly, the negative growth effects of the PathoCom were greatly reduced in the MixedCom experiment. Plants infected with MixedCom grew to a similar extent as the control, with the exception of the genotype Ey15-2, which continued to suffer a substantial weight reduction when infected by the mixed community (**Figure 2**; mean difference to Control = −48.5 mg, [−74.8, −22.6] at 95% confidence interval). Nonetheless, this reduction was less than that caused on Ey15-2 by PathoCom. Hence, co-colonization of pathogenic *Pseudomonas* with commensals led to enhanced growth, while the magnitude was host-genotype dependent.

These results support the role of ATUE5 strains as pathogenic, and provide additional evidence for protection against ATUE5 by commensal *Pseudomonas* strains that coexist with ATUE5 in nature. Next, we wanted to learn whether and how changes in bacterial abundance or shifts in *Pseudomonas* community composition led to differential impacts on growth of the infected plants.

### Differences in bacterial load and impact per a given load of pathogenic and commensal *Pseudomonas*

We hypothesized that the total cumulative load of all barcoded isolates (i.e., regardless of the identity of the colonizing isolates) should be a significant explanatory variable for weight differences among treatments. We based this expectation on the association previously found between abundance in the field and pathogenicity for similar *Pseudomonas* isolates [11].

Contrary to our hypothesis, we found that while the differences in plant weight between treatments were considerable, the bacterial loads of MixedCom and PathoCom were not significantly different from one another (**Figure 3A**). This result implies that plant weight is also a function of bacterial composition, and not load *per se*. In agreement with this inference, the load-weight relationships were found to be treatment-dependent, indicating that weight can be better predicted by load within a treatment than among treatments (difference in expected log-scaled predictive density = −52.9 and in standard error = 9.4 when comparing the model weight ∼ treatment * log_10_(isolates load) + genotype + experiment + error to the same model without the treatment factor, using leave-one-out cross-validation; see Methods).

**Figure 3.**
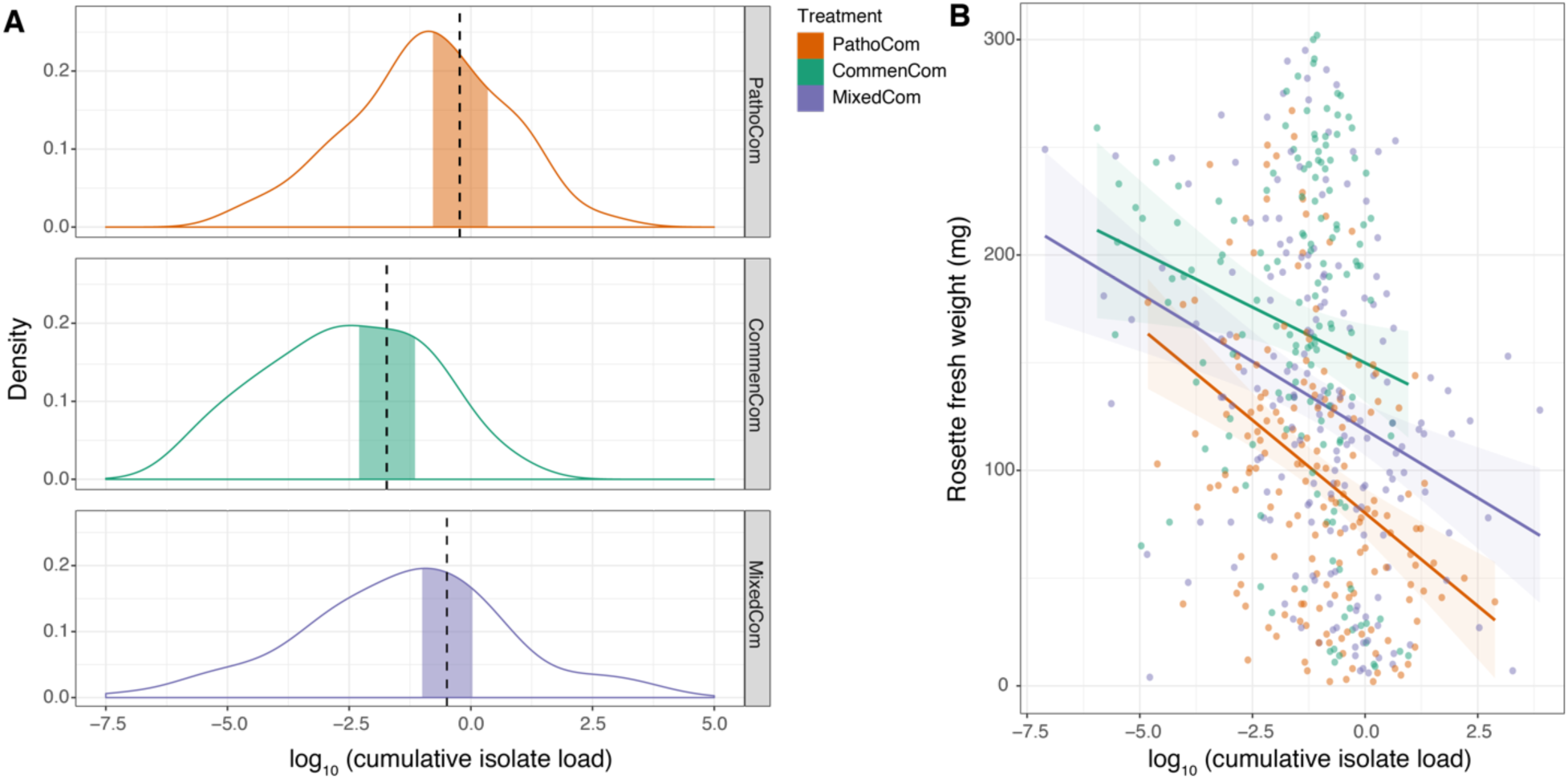
Plants are more tolerant to commensals than pathogens. **A.** Density plot of log_10_(bacterial load) for the three synthetic communities. Vertical dashed lines indicate means, and the shaded areas 95% credible intervals of the fitted parameter, following the model log_10_(bacterial load) ∼ treatment + genotype + experiment + error. **B.** Correlation of log_10_(bacterial load) with rosette fresh weight. Shaded areas indicate 95% confidence intervals of the correlation curve; Bacterial load was defined as the cumulative abundance of all barcoded isolates that constituted a synthetic community. n=170 for PathoCom, n=151 for CommenCom, and n=182 for MixedCom.

Notably, we noticed that the regression slope of PathoCom was more negative than the regression slope of CommenCom, suggesting that ATUE5 isolates had a stronger negative impact on weight per bacterial cell than non-ATUE5 isolates (**Figure 3B**; **Figure S8A**; CommenCom mean effect difference to PathoCom: 12.0 mg [4.4,19.5], at 95% credible interval of the parameter log_10_(isolates load) * treatment). From the reciprocal angle, that of the host, it can be seen that plants were less tolerant to ATUE5 isolates than non-ATUE5 isolates. MixedCom presented a regression slope between the two exclusive synthetic communities, implying that the impact on plant growth resulted from both groups - ATUE5 and non-ATUE5 (MixedCom mean effect difference to PathoCom: 4.8 mg [−1.6,11.8], at 95% credible interval of the parameter log_10_(isolates load) * treatment). Lastly, we observed differential regression slopes between the host genotypes, particularly among Pathocom- and CommenCom-infected hosts, revealing differential tolerance levels to the same *Pseudomonas* isolates (**Figure S8B-C**).

We have described two general differences between pathogenic and commensal *Pseudomonas*: (i) on average pathogens have a greater impact per a given load on plant growth, and (ii) they can reach higher titers in *A. thaliana* leaves. Together, this points to dual effects of pathogens on plant health. In order to explain how commensal non-ATUE5 isolates were able to mitigate the harmful impact of pathogenic ATUE5 in MixedCom, we next addressed the bacterial compositionality in MixedCom-infected hosts.

### Protection by commensal members and host-mediated pathogen suppression

Given that (i) MixedCom-infected plants grew better than PathoCom-infected plants (**Figure 1A**; **Figure S6A**), (ii) there was no considerable difference in total load between PathoCom- and MixedCom-infected plants (**Figure 3A**), and (iii) pathogens were found to cause more damage per cell (**Figure 3B**; **Figure S8A**), we expected commensal members to dominate MixedCom.

Consistent with our expectations, the composition of MixedCom was more similar to CommenCom than PathoCom (**Figure 4A**). We then analyzed the change in bacterial abundance due to the mixture of pathogens and commensals at the isolate level. We compared the absolute abundance of each isolate among the treatments: Pathogenic isolates were compared between PathoCom and MixedCom, and commensals between CommenCom and MixedCom. In general, the abundance of pathogens was significantly lower in MixedCom, while the abundance of commensals was either similar or slightly higher in MixedCom (**Figure 4B**). Thus, the mixture of pathogens and commensals led to pathogen suppression, while commensal load was largely unchanged in MixedCom compared to CommenCom. Thus, non-ATUE5 isolates appear to be more competitive in the MixedCom context than ATUE5 isolates. The abundance change of each isolate in the presence of additional community members was similar among the host genotypes, implying that commensal-pathogen interactions were majorly a general trait, possibly independent of the host (**Figure S9, Table S3**).

**Figure 4.**
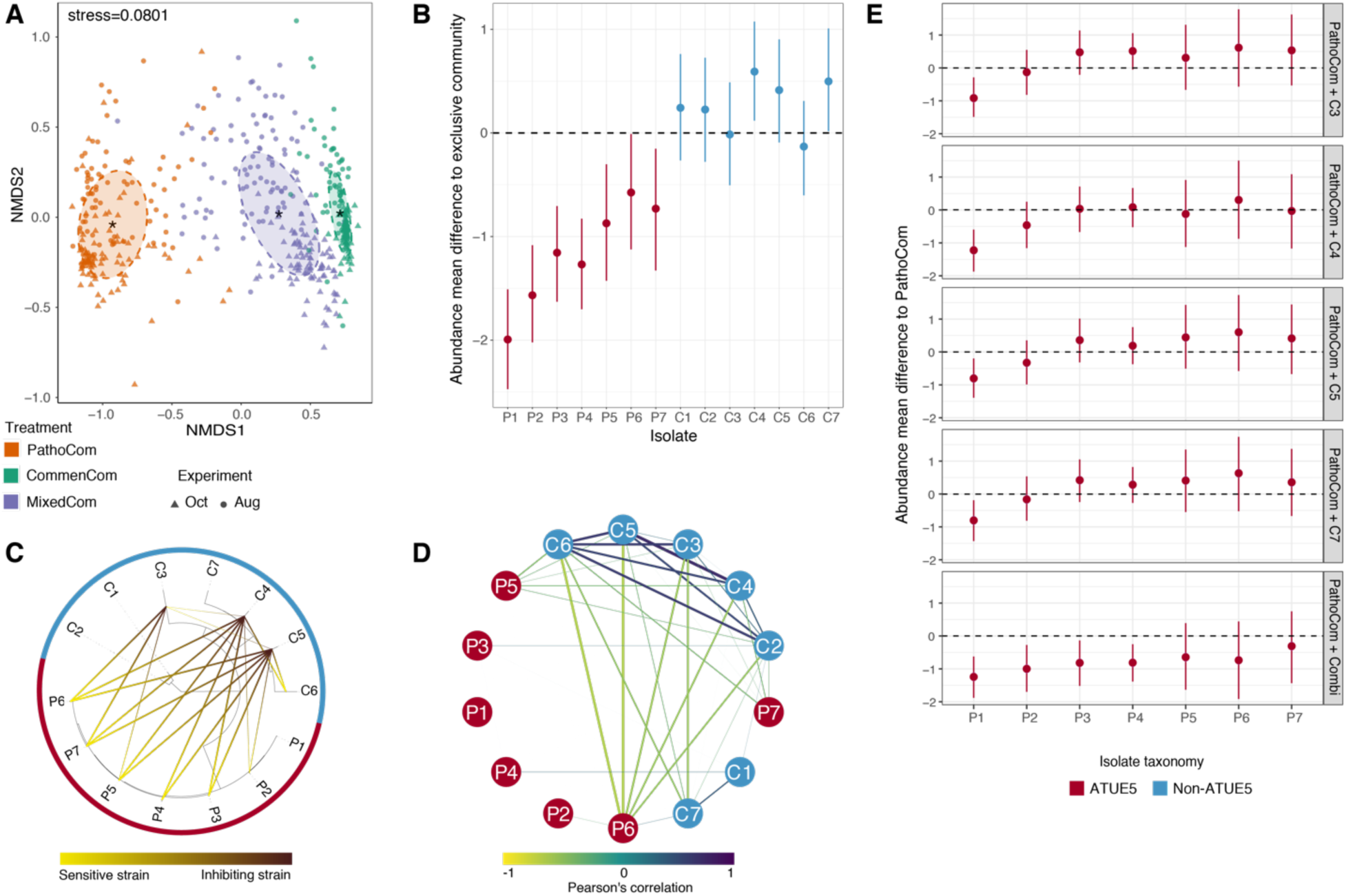
Differential inhibition patterns of pathogens by commensals *in vitro* and *in planta*. **A.** Nonmetric multidimensional scaling (NMDS) based on Bray-Curtis distances between samples infected with the three synthetic communities, across two experiments. The abundance of all 14 barcoded isolates was measured in all communities, including PathoCom and CommenCom, which contained only 7 of the 14 isolates, to account for potential cross contamination and to avoid technical bias. Oct=October, Aug=August. **B.** Abundance change of the 14 barcoded isolates in MixedCom when compared to their exclusive community, in infected plants (i.e. PathoCom for ATUE5, and CommenCom for non-ATUE5). Abundance mean difference was estimated with the model log_10_(isolate load) ∼ treatment * experiment + error, for each individual strain. Thus, the treatment coefficient was estimated per isolate. Dots indicate the medians, and vertical lines 95% credible intervals of the fitted parameter. **C.** Taxonomic representation of the 14 barcoded isolates tested *in vitro* for directional interactions. Ring colors indicate the bacterial isolate classification, ATUE5 or non-ATUE5. Directional inhibitory interactions are indicated from yellow to black. **D.** Correlation network of relative abundances of all 14 barcoded isolates in MixedCom-infected plants. Strengths of negative and positive correlations are indicated from yellow to purple. Boldness of lines is also indicating the strength of correlation, and only correlations > |±0.2| are shown. Node colors indicate the bacterial isolate classification, ATUE5 or non-ATUE5. **E.** *in planta* Abundance change of the seven ATUE5 isolates in non-ATUE5 inclusive treatments, in comparison to PathoCom. Abundance mean difference was estimated with the model log_10_(isolate load) ∼ treatment * experiment + error, for each individual strain. Thus, the treatment coefficient was estimated per isolate. Dots indicate the medians, and vertical lines 95% credible intervals of the fitted parameter. ‘Combi’ - combination of the isolates C3,C4,C5 and C7.

We therefore tested for direct, host-independent interactions among isolates with an *in vitro* growth inhibition assay (Methods). Each of the 14 isolates was examined for growth inhibition against all other isolates, covering all possible combinations of binary interactions. In total, three strains out of the 14 had inhibitory activity; all were non-ATUE5 (**Figure 4C**). Specifically, C4 and

C5 showed the same inhibition pattern: Both inhibited all pathogenic isolates but P1, and both inhibited the same two commensals, C6 and, only weakly, C3. C3 inhibited a total of three ATUE5 isolates: P5, P6 and P7. In summary, the *in vitro* assay provides evidence that among the tested *Pseudomonas*, direct inhibition was a trait unique to commensals, and susceptible bacteria were primarily pathogens. This supports the notion that ATUE5 and non-ATUE5 isolates have divergent competition mechanisms, or at least differ in the strength of the same mechanism.

The *in vitro* results recapitulated the general trend of pathogen inhibition found among treatments *in planta*. Nevertheless, we observed major discrepancies between the two assays. First, P1 was not inhibited by any isolate in the host-free assay (**Figure 4C**), though it was the most inhibited member *in planta*, among the communities (**Figure 4B**). Second, no commensal isolate was inhibited *in plana*, among communities (**Figure 4B**), while two commensals - C3 and C6 - were inhibited *in vitro* (**Figure 4C**). Both could suggest an effect of the host on microbe-microbe interactions. To explore such effects, we analysed all pairwise microbe-microbe abundance correlations within MixedCom-infected hosts. When we used absolute abundances, all pairwise correlations were positive, also in CommenCom and PathoCom (**Figure S10A**), consistent with there being a positive correlation between absolute abundance of individual isolates and total abundance of the entire community (**Figure S11**), i.e., no isolate was less abundant in highly colonized plants than in sparsely colonized plants. It indicates that there does not seem to be active killing of competitors *in planta* in the CommenCom, which is probably not surprising. With relative abundances, however, a clear pattern emerged, with a cluster of commensals that were positively correlated, possibly reflecting mutual growth promotion, and several commensal strains being negatively correlated with both P6 and C7, possibly reflecting unidirectional growth inhibition (**Figure 4D**). We did not observe the same correlations within CommenCom among commensals and within PathoCom among pathogens as we did for either subgroup in MixedCom, reflecting higher-order interactions (**Figure S10B**).

The *in planta* patterns, measured in complex communities, did not fully recapitulate what we had observed *in vitro*, with pairwise interactions. We therefore investigated individual commensal isolates for their ability to suppress pathogens *in planta*, and also tested the entourage effect. We focused on the three commensals C3, C4 and C5, which had directly inhibited pathogens *in vitro*, and C7, which had not shown any inhibition activity *in vitro*, as control. We infected plants with mixtures of PathoCom and each of the four individual commensals, as well as PathoCom mixed with all four commensals. Since pathogen inhibition seemed to be independent of the host genotype, we arbitrarily chose HE-1. Regardless of the commensal isolate, only P1 was significantly suppressed in all commensal-including treatments (**Figure 4E**), with P2,P3 and P4 being substantially inhibited only by the mixture of all four commensals. Together with the lack of meaningful inhibitory difference between individual commensals, this indicates that pathogen inhibition was either a function of commensal dose, or a result of interaction among commensals.

An important finding was that four commensal strains had much more similar inhibitory activity *in planta* than *in vitro*, and that the combined action was greater than the individual effects. Together, this suggested that the host contributes to the observed interactions between commensal and pathogenic *Pseudomonas*. To begin to investigate this possibility, we studied potential host immune responses with RNA sequencing.

### Defensive response elicited by non-ATUE5 inferred from host transcriptome changes

For the RNA-seq experiment, we treated plants of the genotype Lu3-30 with the three synthetic communities, and also used a bacteria-free control treatment. We sampled the treated plants at three and four days after infection (dpi), thus increasing the ability to pinpoint differentially expressed genes (DEGs) between treatments that are not highly time-specific. Exploratory analysis indicated that the two time points behaved similarly, and they were combined for further in-depth analysis.

We first looked at DEGs in a comparison between infected plants and control; with PathoCom, there were only 14 DEGs, with CommenCom 1,112 DEGs, and MixedCom 1,949 DEGs, suggesting that the CommenCom isolates, which are also present in the MixedCom, elicited a host stronger response than the PathoCom members. Furthermore, the high number of DEGs in MixedCom - higher than both PathoCom and CommenCom together - suggest a synergistic response derived from inclusion of both PathoCom and CommenCom members. Alternatively, this could also be a consequence of the higher initial inoculum in the 14-member MixedCom than either the 7-member PathoCom or 7-member CommenCom, or a combination of the two effects (**Figure 5A-B; Figure S12**).

**Figure 5.**
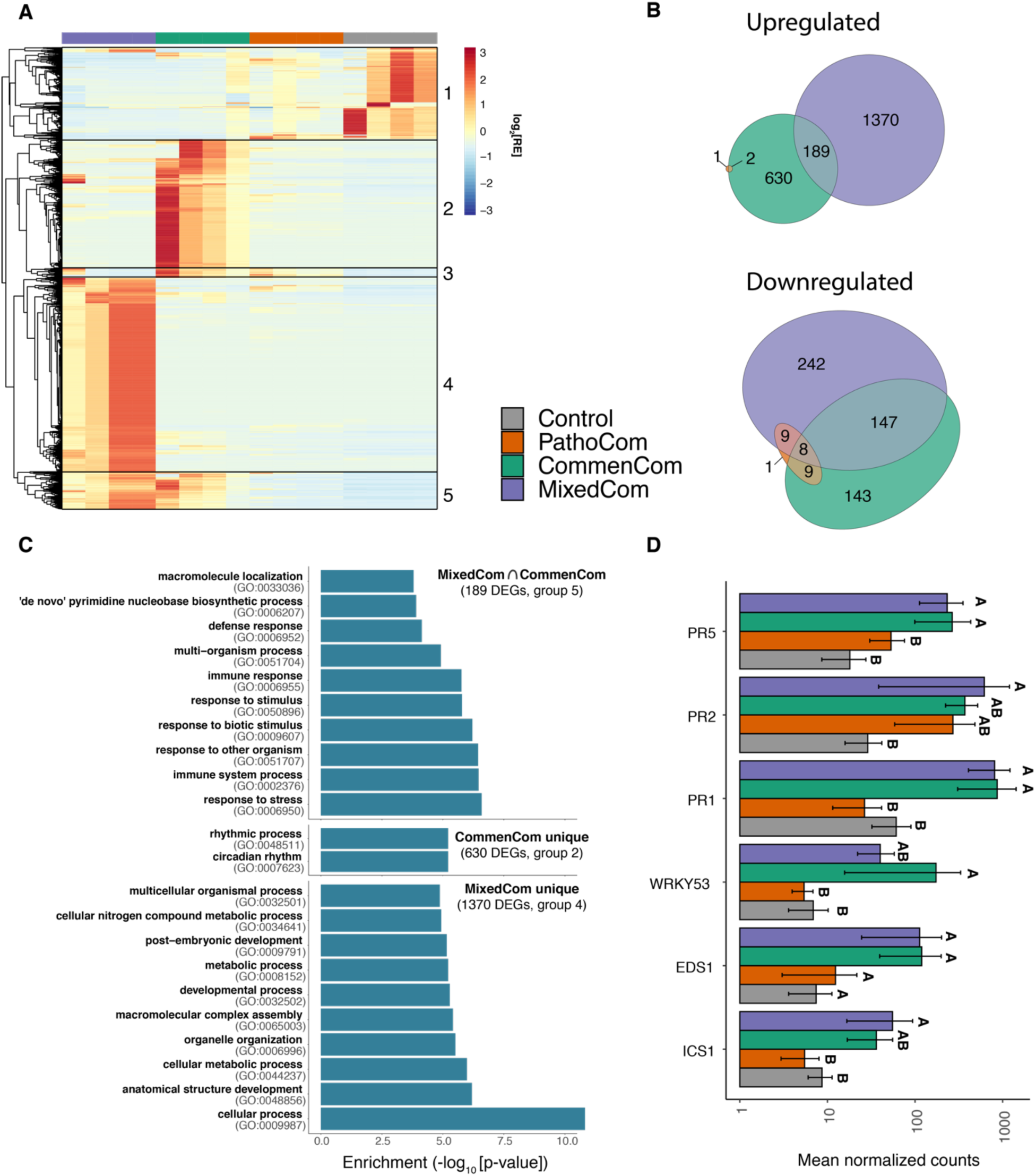
Only commensal members elicit a host-defensive response. **A.** Relative expression (RE) pattern of 2,727 differentially expressed genes (DEGs) found in at least one of the comparisons of CommenCom, PathoCom and MixedCom with control. DEGs were hierarchically clustered. **B.** Euler diagram of DEGs in PathoCom-, CommenCom- and MixedCom-treated plants, compared with control (log2[FC] > |±1|; FDR < 0.05; two-tailed Student’s *t*-test followed by Benjamini-Hochberg correction). **C.** Overrepresented GO terms in upregulated DEG subsets: CommenCom and MixedCom intersection (189 DEGs), CommenCom unique (630 DEGs) and MixedCom unique (1,370 DEGs). Only the top ten non-redundant GO terms are presented; for the full lists of overrepresented GO terms and expression data, see Table S4 and Supplementary Data 1. **D.** Expression values of six defense marker-genes. Mean ± SEM. Groups sharing the same letter are not significantly different (Tukey-adjusted, P>0.05); n=4.

The genes induced by the MixedCom fell into two classes: Group 5 (**Figure 5A-B**) was also induced, albeit more weakly, by the CommenCom, but not induced by the PathoCom. This group was overrepresented for non-redundant gene ontology (GO) categories linked to defense (**Figure 5C**) and most likely explains the protective effects of commensals in the MixedCom. Specifically, among the top ten enriched GO categories in the shared MixedCom and CommenCom set, eight relate to immune response or response to another organism (‘defense response’, ‘multi−organism process’, ‘immune response’, ‘response to stimulus’, ‘response to biotic stimulus’, ‘response to other organism’, ‘immune system process’, ‘response to stress’)(**Figure 5C**).

Group 4 was only induced in MixedCom, either indicating synergism between commensals and pathogens, or being a consequence of the higher initial inoculum. This group included a small number of redundant GO categories indicative of defense, such ‘salicylic acid mediated signaling pathway’, ‘multi-organism process’, ‘response to other organism’ and ‘response to biotic stimulus’ (**Table S4**). Moreover, the MixedCom response cannot simply be explained by synergistic effects or commensals suppressing pathogen effects, since there was a prominent class, Group 2, which included genes that were induced in the CommenCom, but to a much lesser extent in the PathoCom or MixedCom. From their annotation, it was unclear how they can be linked to infection (**Figure 5C**).

About 500 genes (Group 1) that were downregulated by all bacterial communities are unlikely to contain candidates for commensal protection (**Figure 5A**).

Cumulatively, these results imply that the CommenCom members elicited a defensive response in the host regardless of PathoCom members, while the mixture of both led to additional responses. To better understand if selective suppression of ATUE5 in MixedCom infections may have resulted from the recognition of both non-ATUE5 and ATUE5 (reflected by a unique MixedCom set of DEGs) or solely non-ATUE5 (a set of DEGs shared by MixedCom and CommenCom), we examined the expression of key genes related to the salicylic acid (SA) pathway and downstream immune responses. Activation of the SA pathway was previously related to increased fitness of *A. thaliana* in the presence of wild bacterial pathogens, a phenomenon which was attributed to an increased systemic acquired resistance (SAR) [28]. We observed a general trend of higher expression in MixedCom- and CommenCom-infected hosts for several such genes (**Figure 5D**). Examples are *PR1* and *PR5*, marker genes for SAR and resistance execution. Therefore, according to the marker genes we tested, non-ATUE5 elicited a defensive response in the host, regardless of ATUE5 presence.

We conclude that the expression profile of non-ATUE5 infected Lu3-30 plants suggests an increased defensive status, supporting our hypothesis regarding host-mediated ATUE5 suppression. We note, however, that ATUE5 suppression was not associated with full plant protection (thus control-like weight levels) in all plant genotypes. One, Ey15-2, was only partially protected by MixedCom (**Figure 2**), despite levels of pathogen inhibition being not very different from other host genotypes (**Figure S9**).

### Lack of protection in the genotype Ey15-2 explained by a single pathogenic isolate

The fact that Ey15-2 was only partially protected by MixedCom (**Figure 2**), manifest the importance of the host genotype in plant-microbe-microbe interactions, and reflecting dynamics between microbes and plants in wild populations. We wanted to reveal the cause for this differential interaction.

Our first aim was to rank compositional variables in MixedCom according to their impact on plant weight, regardless of host genotype. Next, we asked whether any of the top-ranked variables could explain the lack of protection in Ey15-2. With Random Forest analysis, we estimated the weight-predictive power of all individual isolates in MixedCom, as well as three cumulative variables: Total bacterial abundance, total ATUE5 abundance, and total non-ATUE5 abundance. We found that the best weight-predictive variable was the abundance of pathogenic isolate P6, followed by total bacterial load and total ATUE5 load - which were probably confounded by the abundance of P6 (**Figure 6A**). In agreement, P6 was the dominant ATUE5 in MixedCom (**Figure 6B, Figure S13A**). We thus hypothesized that the residual pathogenicity in MixedCom-infected Ey15-2 was caused by P6.

**Figure 6.**
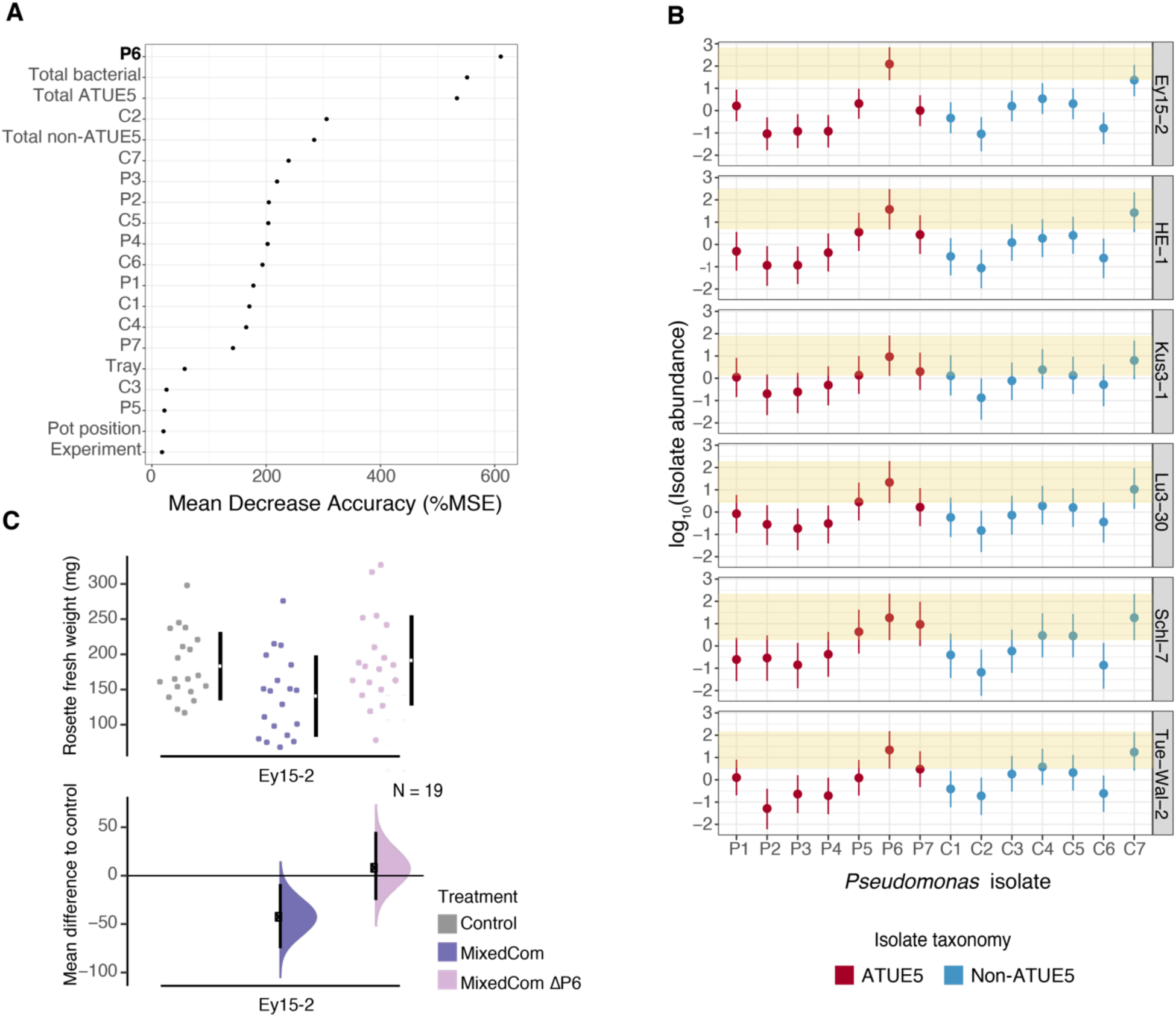
The effect of the isolate P6 on weight in MixedCom-infected hosts, and particularly on the host Ey15-2. **A.** Relative importance (mean decrease accuracy) of 20 examined variables in weight prediction of MixedCom-infected hosts, as determined by Random Forest analysis. The best predictor was abundance of isolate P6. ‘Total Bacterial’, ‘Total ATUE5’ and ‘Total non-ATU5’ are the cumulative abundances of the 14 isolates, 7 ATUE5 isolates, and 7 non-ATUE5 isolates, respectively. **B.** Abundance of P6 compared with the other 13 barcoded isolates in MixedCom-infected hosts, across the six *A. thaliana* genotypes used in this study. Dots indicate the medians, and vertical lines 95% credible intervals of the fitted parameter, following the model log_10_(isolate load) ∼ isolate * experiment + error. Each genotype was analyzed individually, thus the model was utilized for each genotype separately. Shaded area denotes the 95% credible intervals for the isolate P6. **C.** Fresh rosette weight of Ey15-2 plants treated with Control, MixedCom and MixedCom without P6 (MixedCom ΔP6). Fresh rosette weight was measured 12 dpi. The top panel presents the raw data, with the breaks in the vertical black lines denoting the mean value of each group, and the vertical lines themselves indicating standard deviation. The lower panel presents the mean difference to control, plotted as bootstrap sampling [26,27], indicating the distribution of effect size that is compatible with the data. 95% confidence intervals are indicated by the black vertical bars.

Although P6 grew best in Ey15-2, the difference to most other genotypes was not significant (**Figure S13B**). However, P6 was particularly dominant in Ey15-2 (**Figure 6B**).

Given that pathogen load in Ey15-2 was driven to a substantial extent by P6, we assumed that this isolate had a stronger impact on the weight of this genotype than in others. We experimentally validated that removal of P6 restored protection, when Ey15-2 was infected with MixedCom (**Figure 6C**).

Collectively, these results reveal the outcome of direct host-microbe interactions in the context of multiple microbes. Furthermore, they illustrate how plant genotype affects colonization by microbes, and how this may lead to plant health outcomes.

## Discussion

In this work, we aimed to understand how complex interactions between closely related *Pseudomonas* strains affect plant health, considering host-microbe, microbe-microbe and host-microbe-microbe relationships. Not surprisingly, we found that genetics mattered at all levels: membership of *Pseudomonas* strain in commensal or pathogenic clade, genetic variation within each *Pseudomonas* clade, and genetic diversity among *A. thaliana* host strains. Commensal *Pseudomonas* can protect *A. thaliana* from the effects of pathogenic *Pseudomonas* by reducing their proliferation within the plant. However, although this was a general phenomenon, one *A. thaliana* genotype was only partially protected, and this was due to this genotype being particularly susceptible to a specific *Pseudomonas* pathogen. Together, this demonstrates how the host environment can affect microbe-microbe interactions.

The importance of protective interactions for plant health has been demonstrated in both agricultural and wild contexts [1,21,29]. Our results reveal the extreme specificity of these interactions, with closely related pathogenic isolates interacting differently with protective strains We found that upon co-infection with a mixture of pathogens and commensals, pathogens were preferentially suppressed. Perhaps our most important finding was that different plant responses induced by commensals, pathogens and mixed communities. Specifically, commensals, but not pathogens induced transcriptome signatures of defense, and these changes further enhanced in the presence of pathogens. In addition, there were sets of genes that were no longer induced when plants were infected by the mixed community rather than only commensals, as well as sets of genes specifically induced only by the mixed community. This suggests not only that microbe-microbe interactions alter the plant response, but also that these altered plant responses are causal for the differential proliferation of commensals and pathogens in plants affected with mixed communities. These findings support the hypothesis that the complex interplay between the plant immune system and the microbiota goes beyond the individual plant-pathogen interactions, eventually leading to microbial homeostasis [30]. The exact mechanism behind the synergistic effect we describe must still be investigated, though known cases of host-dependent protective interactions provide plausible explanations. For example, early exposure to harmless rhizosphere microbes can prime the plant to suppress at a later time point a broad range of pathogens even in distal tissues, a phenomenon known as induced systemic resistance (ISR) [28].

Another strength of our study is that we used naturally co-occurring biological material, namely strains of *A. thaliana* host and *Pseudomonas* bacteria that had been isolated from a single geographic area. Our results help to explain why the *Pseudomonas* pathogens used here, which are lethal in mono-associations, seem to cause only limited disease in the field [11], namely their effects being modified by other microbes, including other *Pseudomonas* strains.

A limitation of the current study was that we examined only a few commensal isolates, and tested them mostly in complex mixtures. A next logical step will be to test the protective effects of individual commensal *Pseudomonas* strains from the local Tübingen [11] collection, to explore (i) how common protection by commensal *Pseudomonas* is, (ii) how much it depends on the genotype of the pathogen, and (iii) what the genes are that support protection.

We used pathogenic isolates that share over 99% of their 16S rDNA signature, and are highly similar in their core genome [11]. Nonetheless, we found functional differences, relating to both host-microbe and microbe-microbe interactions, exemplified by an individual pathogenic *Pseudomonas* isolate that both dominated the mixed synthetic communities, and that caused a lack of protection in one host genotype. In agreement, Karasov and colleagues [11] had already found that members of this clade of *Pseudomonas* differ substantially in their ability to cause disease in mono-associations.

Friedman and colleagues [31] accurately predicted microbial community structures in the form of trios based on information about pairwise interactions. How easily, however, higher-order communities can be predicted from pairwise interactions, remains to be seen, although recent statistical advances are promising [32,33]. The genome-barcoding method we developed allows strain-level tracking, and thus can be implemented to understand multistrain community assembly. However, in its current format, it is limited to low-throughput studies, mainly due to the cumbersome cloning and transformation serial process. An alternative is presented by high-throughput experiments that combine whole-genome sequencing with statistical reconstitution of known haplotypes [34,35], and which could be employed to study the dynamics of more complex communities. A growing body of literature is revealing effects that can only be found by the ensemble of relationships. For example, in inflammatory bowel disease [36] disease has been linked to changes in microbial community structure rather than to an individual microbe. Another example is provided by plant beneficial consortia, in which only microbial mixtures, but not any single strain triggered pathogen suppression [37,38].

Further advancements in understanding of the plant-microbe-microbe complex in the light of plant health can improve our agriculture practices, allowing the development of more sustainable plant protection methods [39–41].

## Methods

### Plant material

The plant genotypes HE-1, Lu3-30, Kus3-1, Schl-7, Ey15-2 and Tue-Wal2 were used in this study, all originally collected from around Tuebingen, Germany. More details, including stock numbers, can be found in Table S5. Seeds were sterilized by overnight incubation at −80°C, followed by ethanol washes (shake seeds for 5-15min in solution containing 75% EtOH and 0.5% Triton-X-100, and then wash seeds with 95% EtOH and let them dry in a laminar flow hood). Seeds were stratified in the dark at 4°C for 6-8 days prior to planting on potting soil (CL T Topferde; www.einheitserde.de). Plants were grown in 60-pots trays (Herkuplast Kubern, Germany), in which compatible mesh-net pot baskets were inserted, to allow for subsequent relocation of the pots. All plants were grown in short days (8 h of light) at 23°C. Light was applied using Cool White Deluxe fluorescent bulbs, at 125 to 175 μmol m-2 s-1. Relative humidity was set to 65%.

### Barcoding *Pseudomonas* isolates

Excluding the *E. coli* strains that were used for cloning, all 14 bacterial isolates used in this study were classified as *Pseudomonas* and collected from two locations around Tuebingen (Germany) by Karasov and colleagues [11]. Full list, including metadata can be found in Table S1. The procedure of genome-barcoding of the 14 bacterial isolates included random barcodes preparation, cloning the barcodes into pUC18R6KT-mini-Tn7T-Km plasmid and co-transformation of bacteria with the recombinant pUC18R6KT-mini-Tn7T-Km plasmid and pTNS2 helper plasmid (both plasmids from [24]). Preparation of barcodes and the flanking priming sites was done by double stranding two overlapping single strand oligos: One that contains restriction sites, universal priming site, 16 random nucleotides and an overlapping region (Bar1), and another oligo that contains the reverse complement overlapping region, the second universal priming site and restriction sites (Bar2), as illustrated in **Figure S14**; Detailed oligo list in Table S6. The two overlapping single strand oligos were mixed in an equi-molar fashion (5ng each, 2μL in total), together with 0.2 μL Q5 high-fidelity DNA polymerase (New England Biolabs, Ipswich, MA, USA), 1x Q5 5x reaction buffer and 225 μM dNTP in a total reaction volume of 20 μL. The mixture went through a double stranding reaction using a thermocycler (Bio-Rad Laboratories, Hercules, CA, USA), with the following conditions: 95°C for 40 s, 55°C for 60 s and elongation at 72°C for 3 min. The resulting product was cloned into pUC18R6KT-mini-Tn7T-Km plasmid, using the restriction enzymes XhoI and SacI and ligation with T4 DNA-Ligase (Thermo Fisher Scientific, USA). Standard restriction and ligation were conducted, as instructed by the manufacturer protocol. Pir1 competent *E. coli* (Thermo Fisher Scientific, USA) cells were transformed with the ligation product, and subsequently plated on selective Lysogeny broth (LB) agar (1.75%) with 50 ng/mL Kanamycin and 100 ng/mL. Bacterial colonies were validated as successful transformants by PCR with the primers p1 and p2 that are specific for the foreign DNA (detailed oligo list in Table S6). Positive colonies were grown in LB overnight and then used for subsequent plasmid isolation (GeneJET Plasmid Miniprep Kit; Thermo Fisher Scientific, USA). About 150 pUC18R6KT-mini-Tn7T-Km recombinant plasmids were stored at −4°C, each is expected to contain a unique barcode. Sanger sequencing was conducted on a subset of the plasmid library using the primer p1, to validate their barcodes sequence. 14 validated barcodes-inclusive plasmids were randomly selected for the barcoding of the 14 isolates, and these were used for co-transformation together with the plasmid pTNS2 to genome-barcode the selected 14 *Pseudomonas* isolates, as described in [24]. Briefly, *Pseudomonas* strains were grown overnight in LB, pelleted and washed with 300mM sucrose solution to create electrocompetent cells, and were finally electroporated with the recombinant pUC18R6KT-mini-Tn7T-Km (barcodes inclusive) and pTNS2 in a ratio of 1:1. Transformed *Pseudomonas* isolates were grown on selective LB-agar media with 30 mg/mL Kanamycin, and colonies were validated by PCR with the primers p1 and p2 (detailed oligo list in Table S6; Gel electrophoresis results in Figure S2A). Positive colonies were grown in LB with 30 mg/mL Kanamycin overnight, and one portion was stored at −80°C in 25% glycerol, while the other portion was used for DNA extraction (Puregene DNA extraction kit; Invitrogen, USA), followed by Sanger sequencing to validate the barcodes sequences (sequences detailed in Table S1).

### Barcoded and WT isolates growth comparison assay

To compare the growth of the 14 barcoded bacteria with their respective WT, both barcoded and WT isolates were grown overnight in Lysogeny broth (LB) and 10 mg/mL Nitrofurantoin (antibiotic in which all isolated *Pseudomonas* can grow), diluted 1:10 in the following morning and grown for 3 additional hours until they entered log phase. Subsequently, bacteria were pelleted at 3500 g and resuspended in LB to a concentration of OD_600_ = 0.0025, in a 96-wells format plate with a transparent, flat bottom (Greiner Bio One, Austria). Finally, the plate was incubated in a plate reader at 28°C while shaking, for 10 hours (Robot Tecan Infinite M200; Tecan Life Sciences, Switzerland). OD_600_ was measured in one hour intervals.

### Synthetic communities infections and plant sampling

All synthetic communities were prepared as followed: The relevant barcoded isolates were grown overnight in Lysogeny broth (LB) and 30 mg/mL Kanamycin, diluted 1:10 in the following morning and grown for 3 additional hours until they entered log phase, pelleted at 3500 g, resuspended in 10 mM MgSO_4_ and pelleted again at 3500 g to wash residual LB, and resuspended again in 10 mM MgSO_4_ to a concentration of OD_600_ = 0.2, creating a stock solution per isolate for subsequent mixtures. Next, the relevant barcoded isolates were mixed to a final solution with a concentration of OD_600_ = 0.0143 per isolate. Thus, the total concentration per synthetic community was OD_600_ = (0.0143 * isolates number), e.g. PathoCom and CommenCom which comprised 7 isolates had a total concentration of OD_600_ = ∼0.1, and MixedCom which comprised 14 isolates had a total concentration of OD_600_ = ∼0.2. The prepared volume for any synthetic community was calculated by the function: Final volume = number of plants to infect * 2.5ml. Control treatment was sterile 10 mM MgSO_4_ solution. Heat-killed PathoCom was made by incubating a portion of the living PathoCom in 100°C for two hours. In Murashige and Skoog infections, PathoCom was diluted 1:10, thus infections were done using O.D. 0.01. All solutions with synthetic communities were stored at 4°C overnight, and infections were conducted in the morning of the following day.

Infections in the Murashige and Skoog (MS) sterile system were done as described by Karasov and colleagues [11]. In brief, 12-14 days old plants were infected by drip-inoculating 200 μl of the corresponding treatment onto the whole rosette.

The leaves of soil-grown plants were spray-infected, 21 days post sowing. Spraying was done with an airbrush (BADGER 250-1; Badger Air-Brush Co., USA), and each plant was sprayed on both the abaxial and adaxial side for about 1.5 s each. Plants of the same treatment group were placed together in 60-pots trays (Herkuplast Kubern, Germany), in which compatible mesh-net pot baskets were pre-inserted to allow for subsequent relocation of the pots. After the treatment, the transportable pots were reshuffled in new 60-pots trays to form a full randomized block design, thus each tray contained plants from all treatments, in equal amounts. The randomized trays were covered with a transparent lid to increase humidity (Bigger Greenhouse- 60×40cm; Growshop Greenbud, Germany). Four days post infection, two built-in openings in the lids were opened to allow for better air flow and to limit humidity. Eight days post infection, lids were removed. Twelve days post infection, the rosettes of all treated plants were detached using sterilized scalpel and tweezers, weighted, washed from epiphytes (sterile distilled water, 70% EtOH with 0.1% Triton X-100 and then again with sterile distilled water), dried using sterilized paper towels and sampled in 2ml screw cap tubes prefilled with Garnet sharp particles 1mm (Roth, Germany). Tubes with the sampled plants were flash freezed in liquid N_2_, and stored in −80°C.

### DNA extraction, barcodes PCR and qPCR

Frozen sampled plants were used for DNA extraction suitable for metagenomics, using a protocol that was previously described by karasov and colleagues [11]. Briefly, the samples were subjected to bead-beating in the presence of 1.5% sodium dodecyl sulfate (SDS) and 1 mm garnet rocks, followed by SDS cleanup with 1/3 volume 5 M potassium acetate, and then SPRI beads.

The resulting DNA was used for a two step PCR. The first PCR step amplified the genome-integrated barcodes and added short overhangs, using the primer p3 and the primers p4-p9. The latter are different versions of one primer with frameshifting nucleotides, allowing for better Illumina clustering, and thus sequencing quality, following the method described by [42] (2013; Detailed oligo list in Table S6). Each primer frameshift version was used for a different PCR plate (i.e. 96 samples). The second step primed the overhangs to Illumina adapters for subsequent sequencing, using standard Illumina TruSeq primer sequences. Unique tagging of PCR samples was accomplished by using 96 indexing primers, combined with the six combinations of frameshift primers in the first PCR (as detailed in [42]), allowing demultiplexing of up to 576 samples in one Illumina lane. The first PCR was done in 25 μL reactions containing 0.125 μL TaqI DNA polymerase (Thermo Fisher Scientific, USA), 1x Taq1 10x reaction buffer, 0.08 μM each of forward and reverse primer, 225 μM dNTP and 1.5 μL of the template DNA. The first PCR was run for 94°C for 5 min followed by 10 cycles of 94°C for 30 s, 55°C for 30 s, 72°C for 1 min, and a final 72°C for 5 min. 5 μL of the first PCR product was used in the second PCR with tagged primers including Illumina adapters, in 25 μL containing 0.25 μL Q5 high-fidelity DNA polymerase (New England Biolabs, USA), 1x Q5 5x reaction buffer, 0.08 μM forward and 0.16 μM of reverse (tagging) primer and 200 μM dNTP. The final PCR products were cleaned twice using SPRI beads in a 1:1 bead to sample ratio, and eluted in 15 μL. Samples were combined into one library in an equimolar fashion. Final libraries were cleaned twice using SPRI beads in a 0.6:1 bead to sample ratio to clean the primers from the product, and were finally eluted in half of their original volume. Samples were sequenced by a MiSeq instrument (Illumina), using a 50 bp single-end kit.

In order to estimate the ratio of barcoded *Pseudomonas* to plant chromosomes, two qPCR reactions were conducted - one which is specific to the barcodes, and the other which is plant-specific, targeting the gene GIGANTEA which is normally found in one copy. For barcodes-specific qPCR, the primers p10 and p11 were used, and for plant-specific qPCR the primers p12 and p13 (Table S6). qPCR reactions were done in 10 μL reactions containing x1 Maxima SYBR green qPCR master mix x2, 0.08 μM each of forward and reverse primer and 1 μL of template DNA. All qPCR reactions were run for 94°C for 2 min followed by 94°C for 15 s and 60°C for 1 min in a BioRad CFX384 Real Time System (Biorad, USA) qPCR machine. Reactions were done in triplicates.

### *In vitro* directional suppression assay

All 14 barcoded isolates *in vitro* pairwise interactions were tested following the method described in Helfrich et al. [43], while adjusting the conditions to better fit *Pseudomonas*. Briefly, the 14 barcoded isolates were grown in LB with 30 mg/mL Kanamycin overnight, diluted 1:10 the following morning and regrown. One portion was taken from each isolate after 3 hours (when entering the log phase), diluted to a final concentration of OD_600_ = 0.001 in 15ml LB with 1% agar and immediately poured into a square plate to form a uniform layer containing the test strain. Another portion pelleted at 3500 g, washed from residual LB in 10 mM MgSO_4_, pelleted again at 3500 g in 10 mM MgSO_4_ with half of the original volume. Roughly 1 μL of each strain was printed onto the solidified agar layer containing the putative sensitive strain. Inhibitory interactions were estimated after 1-2 days incubation at 28°C by documenting observable halos. The strength of inhibitions was assessed by the halo size as previously described [43]).

### RNA-sequencing

Plants from the genotype Lu3-30 were infected with Control, PathoCom, CommenCom and MixedCom as described below. Sampling was conducted three and four days post infection, two replicates per treatment in each time point, thus four samples per treatment in total. Plants were sampled using sterilized scalpel and tweezers and were immediately placed in 2ml screw cap tubes prefilled with Garnet sharp particles 1mm (Roth, Germany), flash freezed in liquid N_2_ and stored in −80°C. RNA extraction was conducted on the frozen samples as previously described [44]. Briefly, a guanidine hydrochloride buffer was added to grounded frozen and rosettes, followed by phase separation and sediments removal. Combined with 96% EtOH, the solution was loaded onto a plasmid DNA extraction column (QIAprep Spin Miniprep Kit; Qiagen), and went through several washes before elution of the RNA. mRNA enrichment and sequencing libraries were prepared as previously described [45]. Briefly, mRNA enrichment was done using NEBNext Poly(A) mRNA Magnetic Isolation Module (New England Biolabs, USA), followed by heat fragmentation. Next, First strand synthesis (SuperScript II reverse transcriptase; Thermo Fisher Scientific, USA), and second strand synthesis (DNA polymerase I; New England Biolabs, USA) were conducted, and subsequently end repair (T4 DNA polymerase, Klenow DNA polymerase and T4 Polynucleotide Kinase; New England Biolabs, USA) and A-tailing (Klenow Fragment; New England Biolabs, USA). Nextera-compatible universal adapters [46] were ligated to the product (T4 DNA ligase; New England Biolabs, USA), and i5 and i7 PCR amplification was done (Q5 polymerase; New England Biolabs, USA). Size selection and DNA purification were made using SPRI beads. Samples were sequenced by a HiSeq3000 instrument (Illumina), using a 150 bp paired-end kit.

### Sampling locations map, phylogenetics and isolates abundance in the field

Information about sampling locations of the six *A. thaliana* used in this study was retrieved from the 1001 genome project [25], and *Pseudomonas* sampling locations were retrieved from Karasov and colleagues [11]. The map was plotted using the “ggmap” function of the ggmap R package [47].

Phylogenetic analysis of the 14 selected *Pseudomonas* isolates was done using their core genomes, as they were previously published [11]. Maximum-likelihood phylogenies were constructed with RAxML (v.0.6.0) using GTR+Gamma model [48], and visualization was done by iTOL [49]. The abundance in the field of the selected isolates was estimated by binning similar isolates using a threshold of divergence less than 0.0001 in the core genome. The mean number of substitutions per site taken from the estimated branch length for the core-genome based phylogeny calculated by RAxML. Lastly, the number of binned isolates was divided by the total number of isolates surveyed by Karasov and colleagues [11].

### Growth analysis of WT and barcoded isolates

Growth of both WT and barcoded isolates was analyzed using the function “SummarizeGrowthByPlate” from the Growthcurver R package [50]. The change of barcoded isolates in comparison to their corresponding WT in growth rate, carrying capacity and area under the curve, was calculated by the model: Growth quantity ∼ strain type (i.e. WT/barcoded).

### Plant weight analysis

All rosette fresh weight analyses and visualizations were done using the function “dabest” of the dabestr R package [26,27].

### Combining barcode PCR and qPCR results to estimate bacterial load per isolate

All reads from barcode-PCR sequencing were mapped against a custom barcodes database (Table S1), and a count matrix of all 14 isolates for every plant sample was created. Samples with less than a total of 200 hits were discarded or resequenced (mean=15709.8). Counts were transformed to proportions by dividing the counts of each isolate in the total hits per sample, resulting in relative abundance matrix.

qPCR results were analyzed using the software Bio-Rad CFX Manager with default parameters. Quantification cycle (Cq) values smaller than 32 were discarded, and barcoded bacterial load was determined by the equation 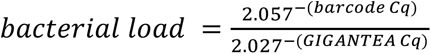. The exponent bases (2.057 and 2.027) were adjusted according to primer efficiency - as determined by a calibration curve derived from a series of dilutions. The relative abundance matrix was factorized by bacterial load (relative abundance multiplied by bacterial load, per isolate) to manifest the ratio of bacterial to plant chromosomes per barcoded isolate.

### Regression analysis

All posterior distributions of focal factors were estimated using the function “stan_glm” in the R package rstanarm ([51] or “lmBF” in the R package BayesFactor ([52]. In both functions, default priors were used. In “stan_glm” default iteration number was used, and in “lmBF” 10,000 iterations were used. In all figures, the median, as well as 2.5% and 97.5% (95% credible intervals) of the posterior distribution were presented for each factor of interest. The exact model for every analysis is presented in the figure legend, as well as the selected references for comparison.

To compare the effect of individual predictors in a model, the full model was compared to a different model, lacking the predictor of interest (e.g. genotype). The comparison was conducted by a leave-one-out cross validation, using the function “loo_compare” in the R package Loo [53]. This Bayesian-based model comparison provides an estimate for the importance of a predictor in explaining the data. Leave-one-out cross validation improves the estimate in comparison to the common Akaike information criterion (AIC) and deviance information criterion (DIC) [53].

### Variance partitioning of microbial community composition

NMDS analyses were conducted using the function “metaMDS” in the R package vegan [54], adjusting dissimilarity index to Bray-Curtis (method = “bray”), number of dimensions to 3 (k=3) and maximal iterations to 200 (trymax=200). Permutational multivariate analysis of variance (PERMANOVA) was conducted using the function “adonis”, and analysis of similarities (ANOSIM) was conducted using the function “anosim” in the R package vegan [54]. Both were adjusted to Bray-Curtis dissimilarity index (method = “bray”) and 2000 permutations (permutations = 2000). Multilevel pairwise comparison using adonis was conducted using the function “pairwise.adonis2” in the R package pairwiseAdonis [55].

### Isolate-isolate interactions network

All pairwise isolate-isolate Pearson correlations were calculated using the function “rcorr” in the R package Hmisc ([56], and visualization was done with Cytoscape 3.7.0 ([57].

### RNA-sequencing analysis

Reads from RNA sequencing were mapped against the *A. thaliana* reference TAIR10 using STAR (v.2.6.0; [58] with default parameters. Transcript counts matrix was done using featureCounts [59], while restricting counts to exons only (-t exon). Differential gene expression (DEG) analysis was conducted using DESeq2 (v.1.22.2; [60]), using the model ‘gene_expression ∼ Treatment + Time_point’. Genes with average counts of less than five were excluded from the analysis. Zero counts were converted to one to allow for the log conversion in unexpressed genes. Genes with log2FoldChange>|±1| and FDR<0.05 (two-tailed Student’s *t*-test followed by Benjamini-Hochberg correction) were defined as DEGs. Euler diagrams were created using the function “euler” in the R package eulerr [61]. Statistically overrepresented GO terms were identified using the BiNGO plugin for Cytoscape [62]. Summarization and the removal of redundant overrepresented GO terms was done with the web server REVIGO [63] to extract the main trends found in the long full output by BiNGO (full list in Table S4).

### Statistical analysis

All statistical analyses were performed using the R environment version 3.5.1, unless mentioned otherwise. Sample sizes were not predetermined using statistical methods.

## Acknowledgments

We thank Alba Gonzalez, Stefan Petschak, Wangsheng Zhu, Wei Yuan, Sonja Kersten, Sergio Latorre, Julian Regalado, Anjar Wibowo, Bridgit Waithaka and Thanvi Srikant for helping with plant sampling. Special thanks to Christine Vogel for her detailed guidance regarding the *in vitro* assay, and for Christoph Ratzke for his input about the manuscript. This work was supported by fellowships from DAAD (O.S.S.), Funding was provided by HFSP long-term fellowships (LT000348/2016-L, T.L.K.; LT000565/2015-L, D.S.L.), an EMBO long-term fellowship (LRTF 1483-2015, T.L.K.) and Alexander von Humboldt Foundation (H.A.), and by DFG SPP DECRyPT and the Max Planck Society.

## Data Availability

RNA sequencing data have been deposited with the European Nucleotide Archive (ENA) under study accession number PRJEB41069.

## Competing Interests

The authors declare no competing interests.

## Author Contributions

O.S.S conceived and designed the research. D.S.L conceived bacterial barcoding and developed the method with O.S.S.. O.S.S performed the experiments and analyzed the results, O.S.S, T.L.K, D.S.L, H.A and D.W. discussed and interpreted the results. O.S.S wrote the first draft and the manuscript was written by O.S.S. and D.W, with input from all authors.

---

**Figure S1.**
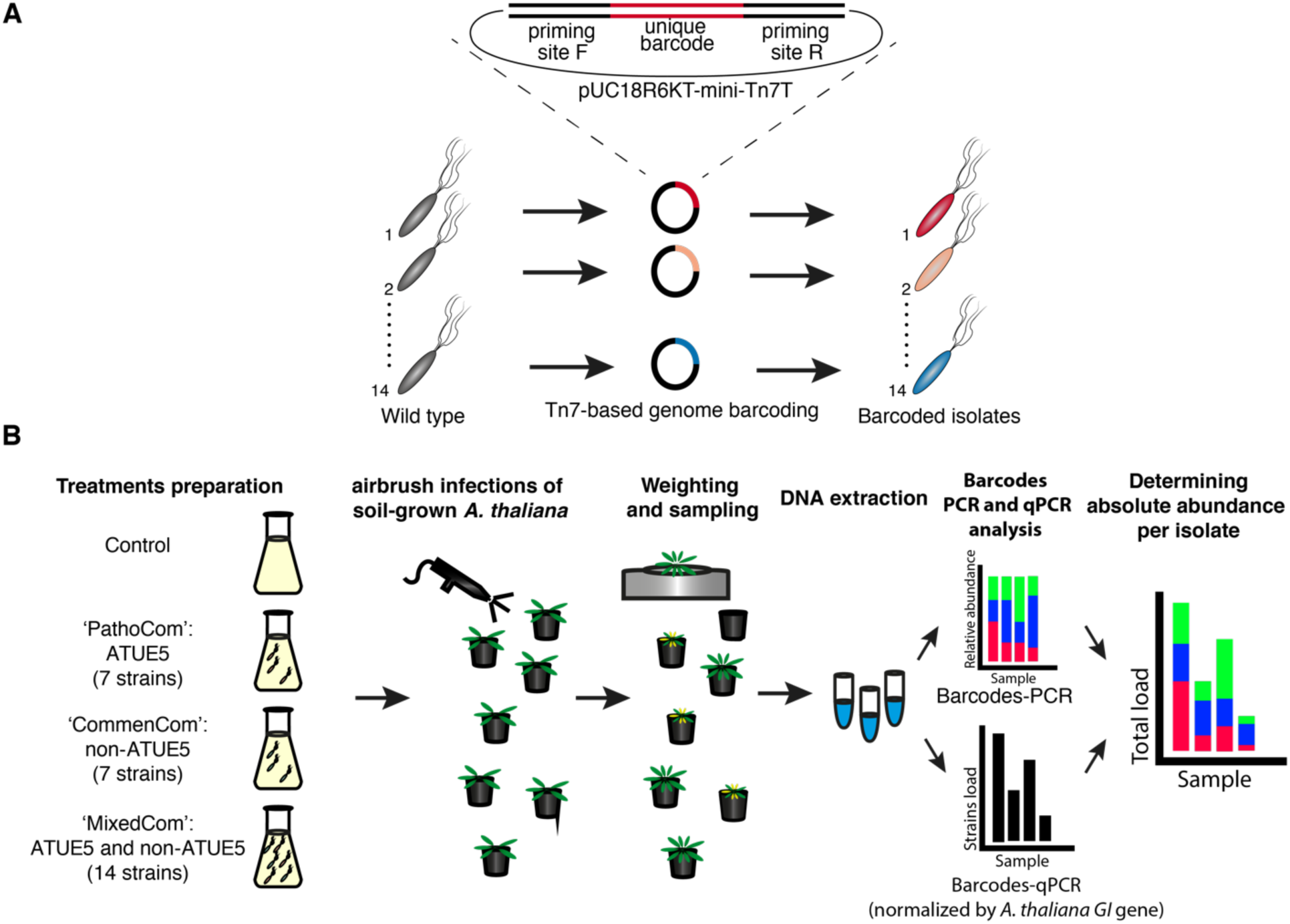
Illustration of (A) bacterial barcoding and (B) experimental design.

**Figure S2.**
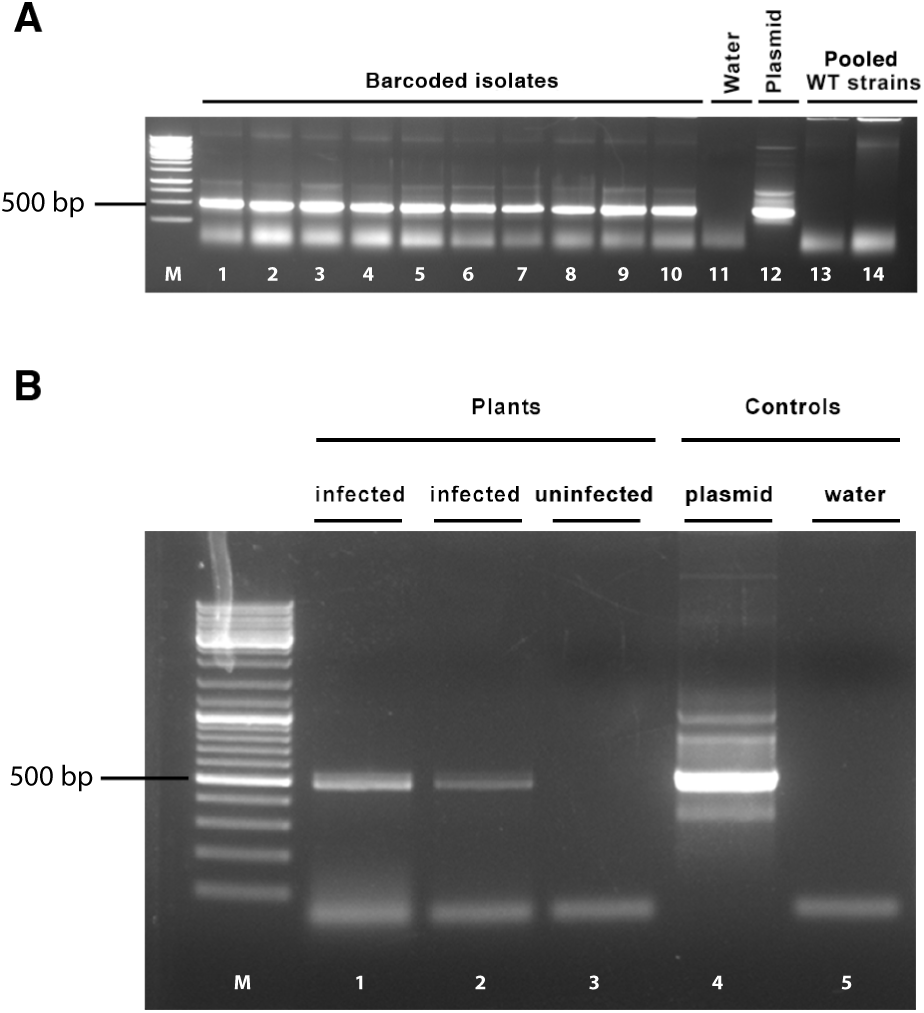
Validation of barcode integration and barcode-PCR specificity by agarose gel electrophoresis of PCR amplified products. **A.** Validation of barcode integration to chosen isolates. Lanes 1–10 used DNA from examined barcoded isolates, lane 11 is water (negative control), lane 12 is the pUC18R6KT-mini-Tn7T plasmid into which a barcode was cloned (positive control), and lanes 13-14 are replicates of the 14 pooled parental (wild-type, WT) isolates. **B.** Validation of barcode-PCR specificity. Lanes 1-2 used DNA from plants infected with the 14 barcoded bacteria, lane 3 from an uninfected plant, lane 4 pUC18R6KT-mini-Tn7T plasmid (positive control), and lane 5 is water (negative control). Both infected and uninfected plants were grown in non-sterile conditions; barcode-specific primer sets yielded expected product sizes of 522 bp. Lane M, DNA size marker. 500 bp marker indicated.

**Figure S3.**
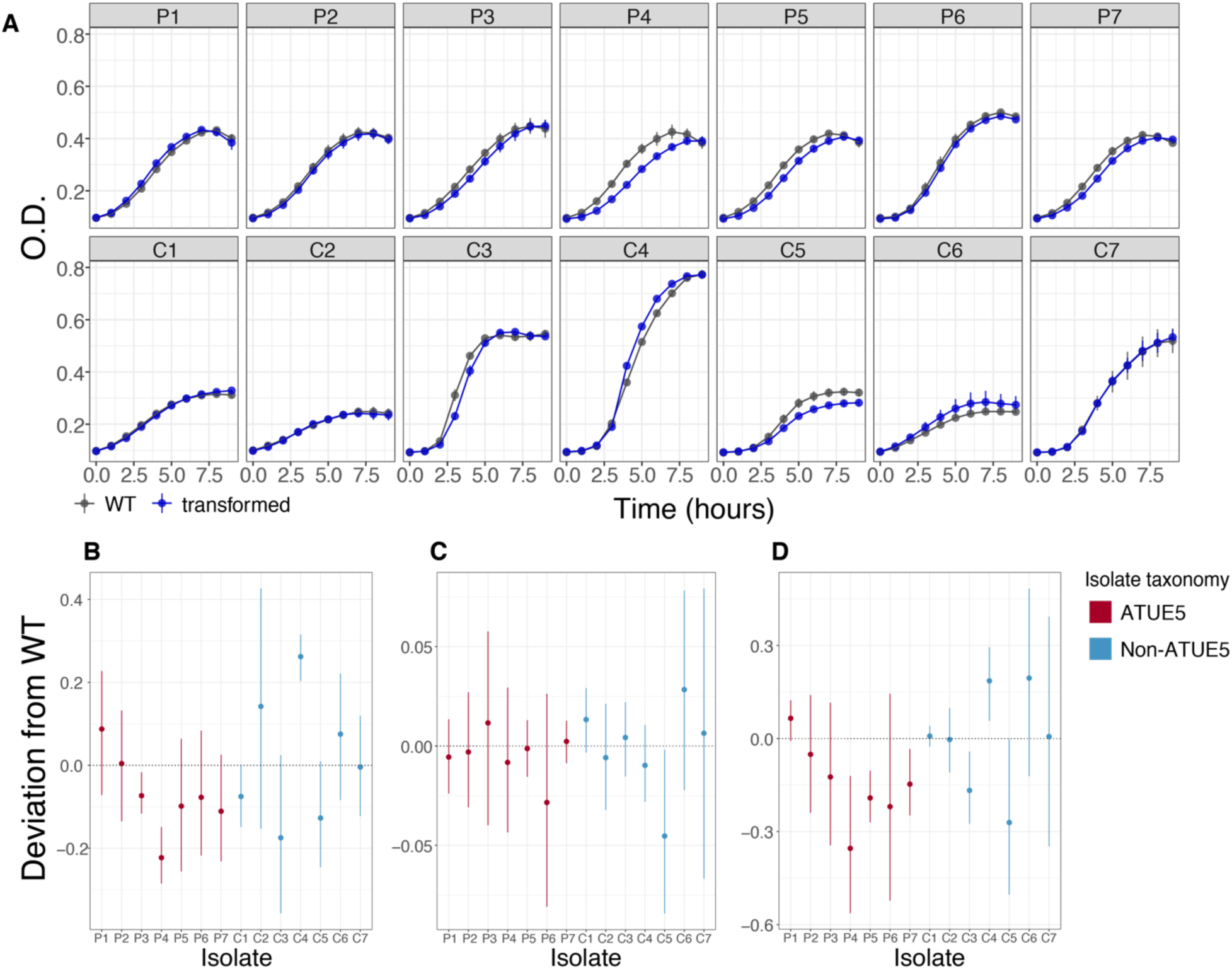
Comparison of growth characteristics between non-barcoded wild-type (WT) isolates and their barcoded derivatives. **A.** Growth curves of the 14 WT parents and their barcoded derivatives in Lysogeny broth (LB) over 10 hours, with OD_600_ recorded hourly. Mean ± SD, n=3. The change of barcoded isolates in comparison to their corresponding parents in growth rate (**B**), carrying capacity (**C**), and area under the curve (**D**) is shown. All three growth parameters were derived from the original growth curves. Dotted line signifies the non-barcoded parental baseline for a given quantity. Mean ± 95% cdl, n=3.

**Figure S4.**
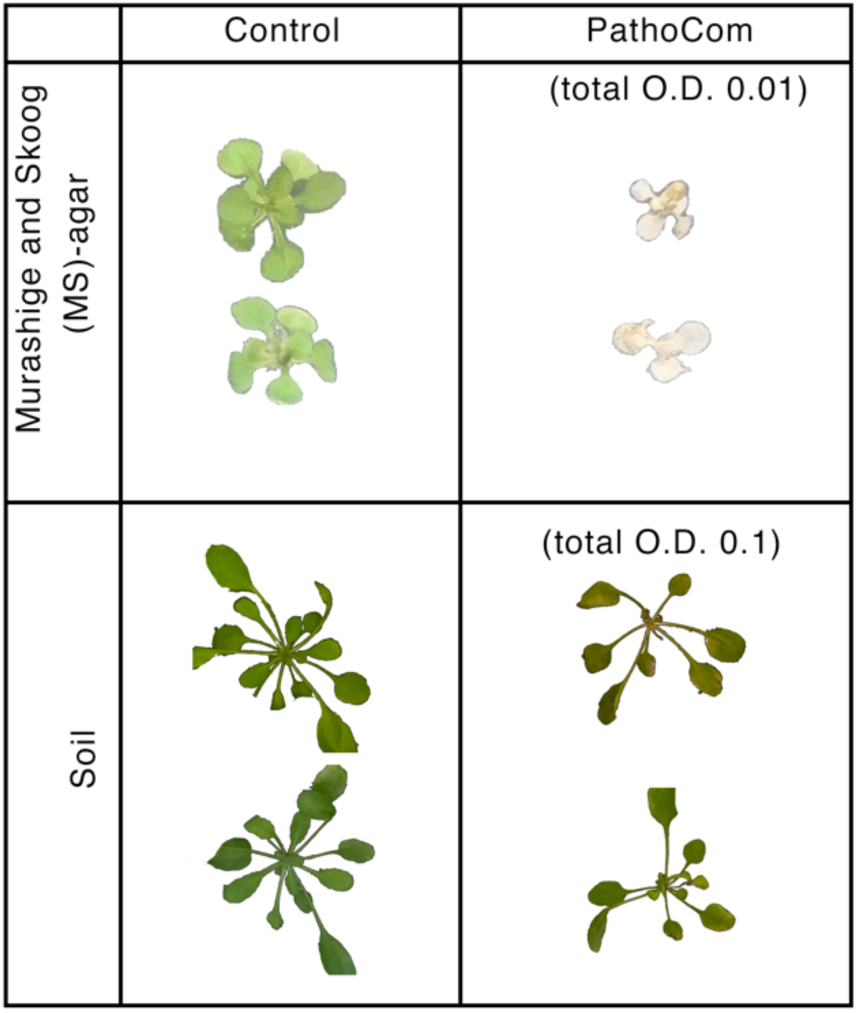
Illustrative photos of control- and PathoCom-treated plants, grown in either MS-agar (sterile) or soil (unsterile). In both systems, the genotype Ey15-2 was used. For the MS-agar system, photos were taken 3-dpi, for the soil system 14-dpi. Sizes of plants are comparable within each system, but not between. Because images in the soil system were taken and parsed by pot automatically by a high-throughput imaging pipeline, some plant images were cropped.

**Table S2.**
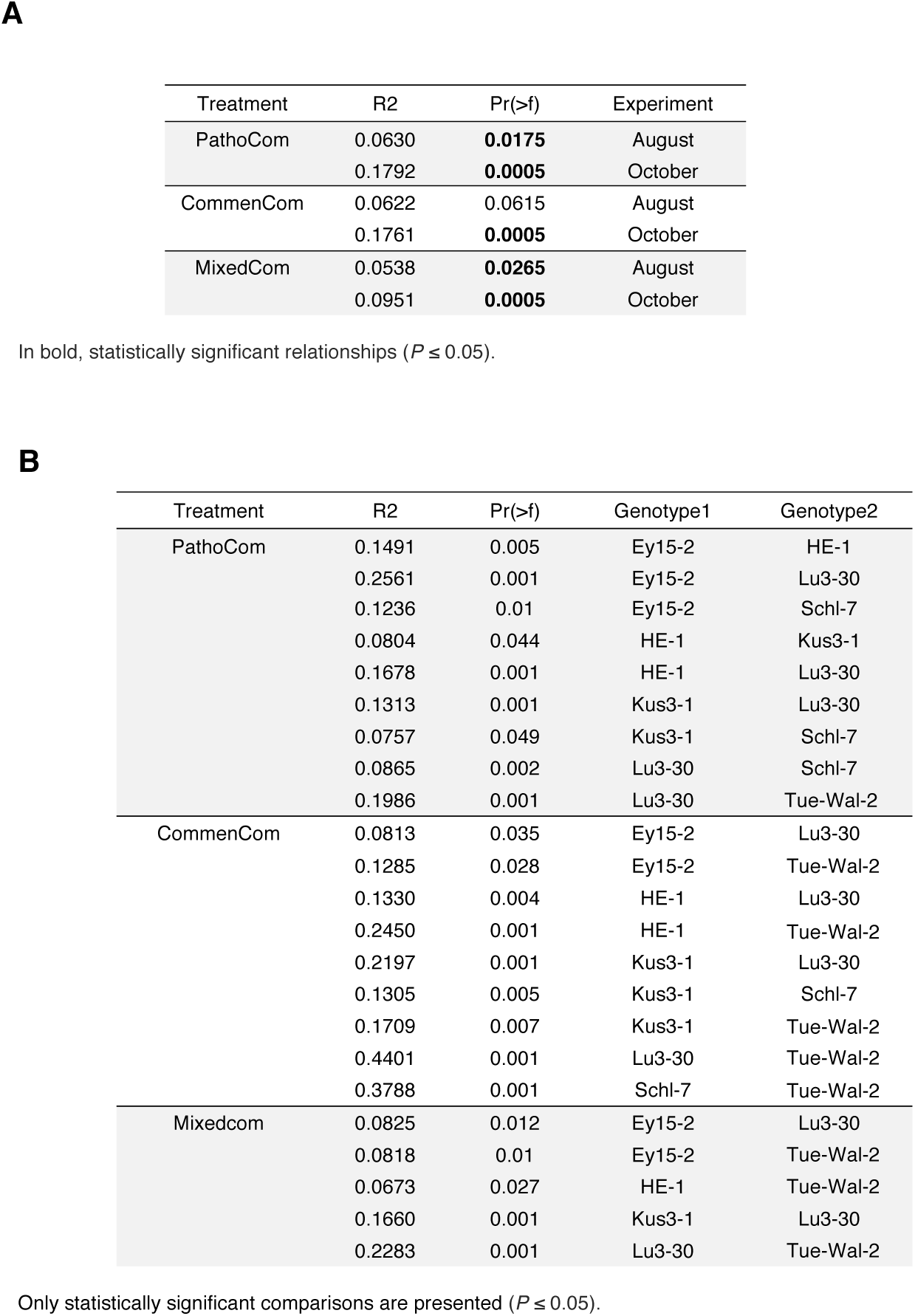
**A.** Analysis of similarities (ANOSIM) based on Bray-Curtis distances for compositions of the 14 barcoded bacteria in treated hosts. The analysis was constrained by the host genotype in each experiment batch (exp) to estimate its effect on the explained variance. **B.** Multilevel pairwise comparison of barcoded bacteria compositions for the different *A. thaliana* genotypes, using adonis based on Bray-Curtis distances. Data derived from one representative experiment (October).

**Figure S5.**
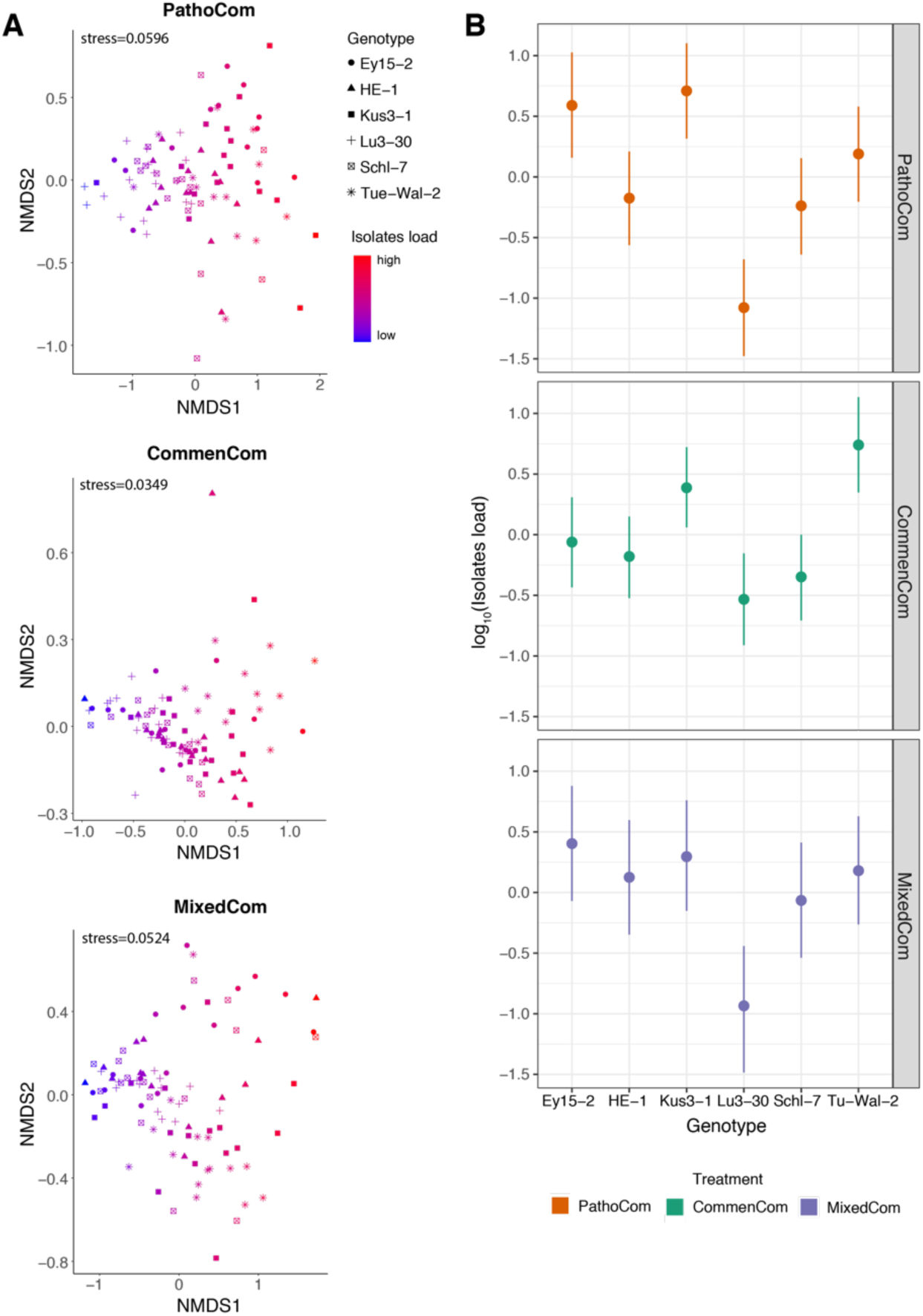
Comparison of composition and load of the 14 barcoded isolates on different *A. thaliana* genotypes. **A.** Nonmetric multidimensional scaling (NMDS) based on Bray-Curtis distances between six *A. thaliana* genotypes, in one representative experiment (October). Each synthetic community was analyzed separately. The abundance of all 14 barcoded isolates was considered, also among PathoCom and CommenCom to account for cross contaminations and technical distortions. Shapes denote the different genotypes, and bacterial load is indicated from blue to red. **B.** Isolate load of the six *A. thaliana* genotypes, among the three synthetic communities. Isolate load was defined as the cumulative abundance of all barcoded isolates that composed a synthetic community. Dots indicate the medians, and vertical lines 95% credible intervals of the fitted parameter, following the model log_10_(isolates load) ∼ genotype + experiment + error.

**Figure S6.**
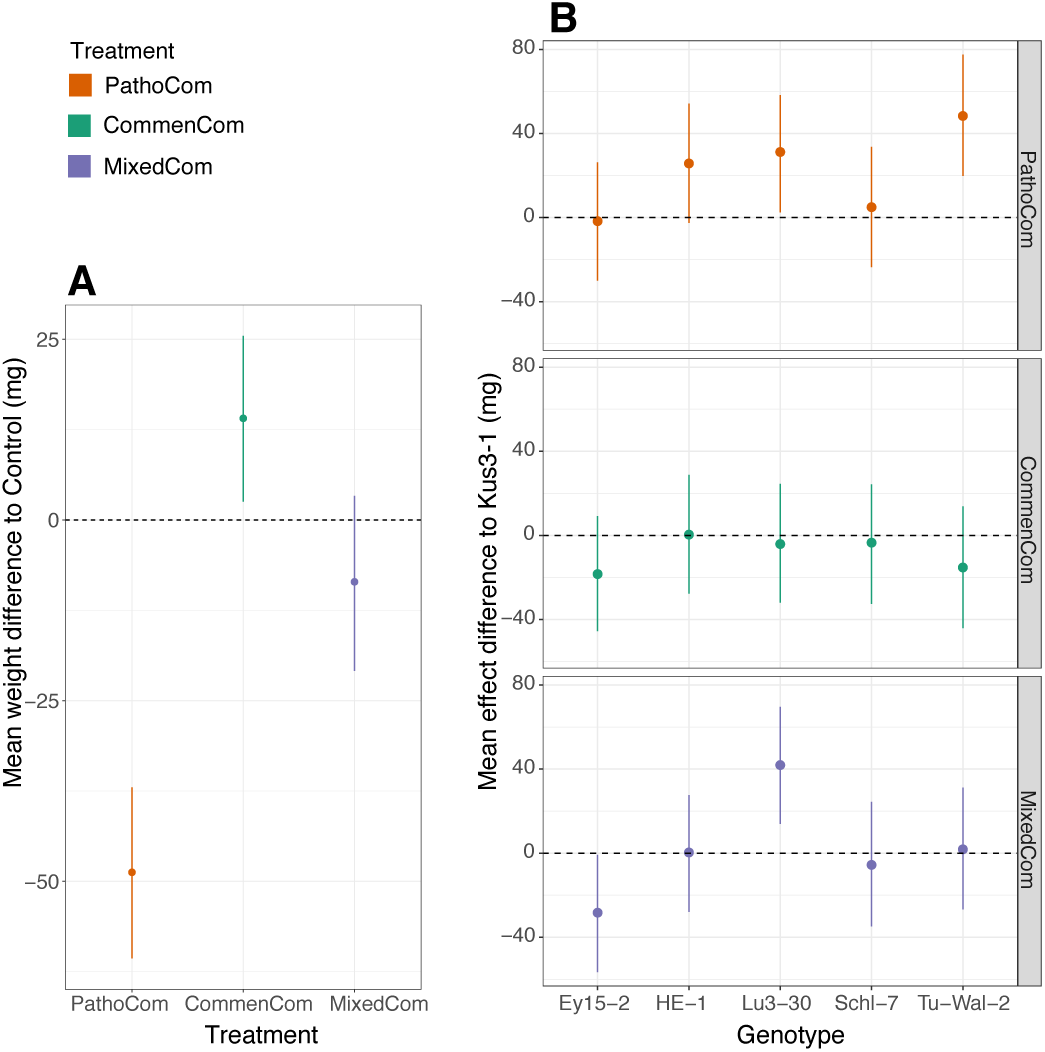
Effects of treatment and treatment-by-genotype on fresh rosette weight. Both effects were assessed using the model weight ∼ treatment * genotype + experiment + error. **A.** Mean weight difference of plants infected with each of the three synthetic communities relative to control - i.e., the treatment coefficients. **B.** Mean treatment effect differences between the six *A. thaliana* genotypes used in this study - i.e., the treatment * genotype coefficients. Kus3-1 was randomly selected as a reference; dots indicate the medians, and vertical lines 95% credible intervals of the fitted parameter.

**Figure S7.**
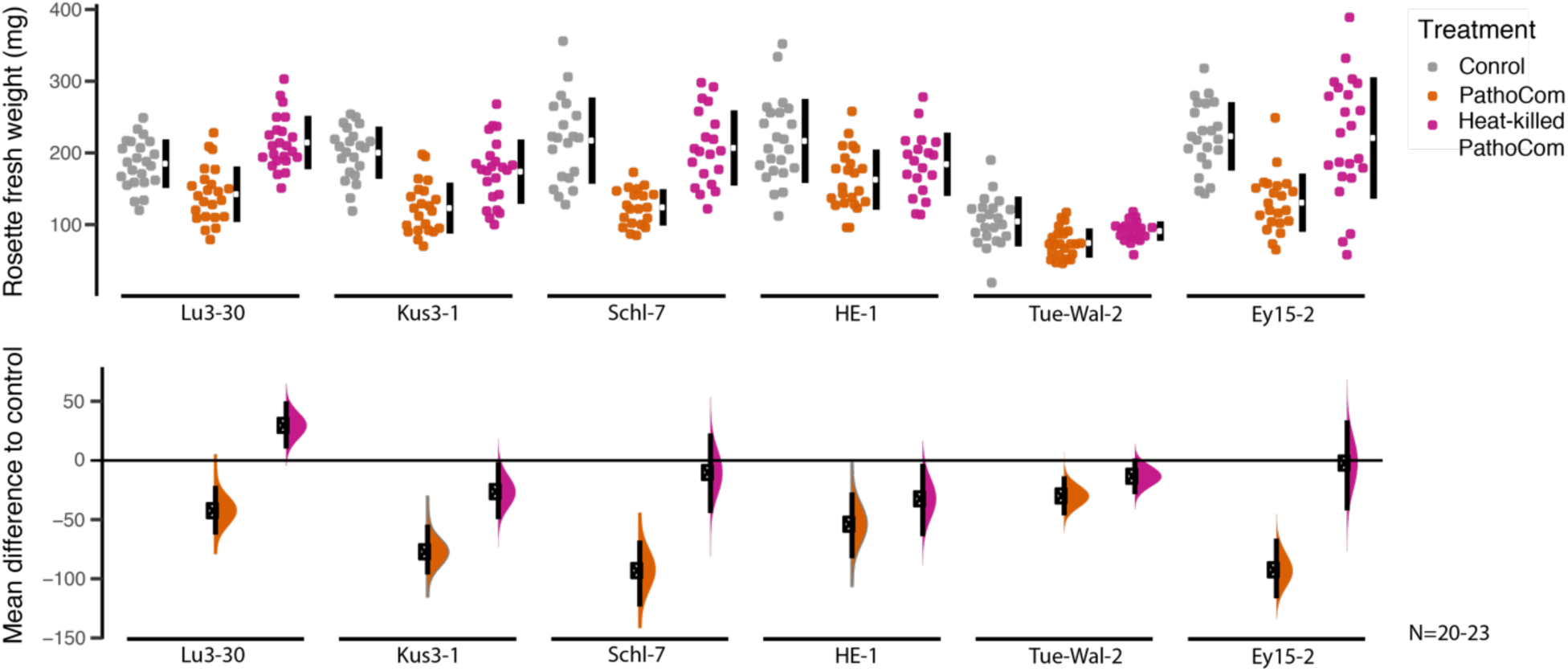
Fresh rosette weight of plants treated with Control, PathoCom or heat-killed PathoCom. Each of the six *A. thaliana* genotypes used in this study was treated with control, PathoCom and heat-killed PathoCom inoculum, and fresh rosette weight was measured 12 dpi. The top panel presents the raw data, the breaks in the vertical black lines denote the mean value of each group, and the vertical lines themselves indicate standard deviation. The lower panel presents the mean differences to control, plotted as bootstrap sampling [26,27], indicating the distribution of effect sizes that are compatible with the data. 95% confidence intervals are indicated by the black vertical bars.

**Figure S8.**
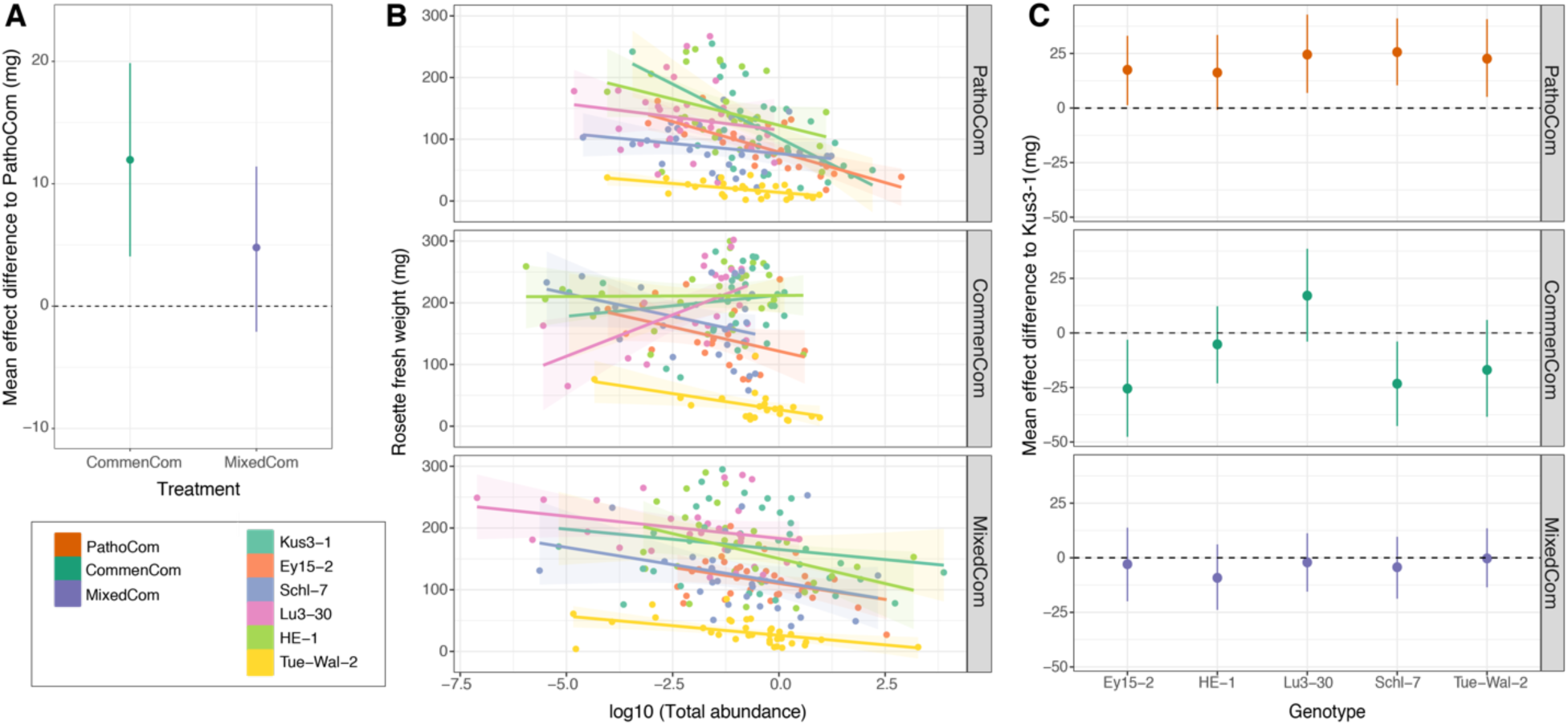
Effect of total load on weight, per treatment and genotype. **A.** Mean slope difference of the three synthetic communities. The slope difference indicates the effect of the treatment on the correlation between weight and isolate load - i.e. treatment * log_10_(cumulative isolate load) - following the model weight ∼ treatment * log_10_(cumulative isolate load) + genotype + experiment + error. PathoCom was selected as a reference. Dots indicate the medians, and vertical lines 95% credible intervals of the fitted parameter. Related to Fig 3B. **B.** Correlation of log_10_(cumulative isolate load) with rosette fresh weight, for each of the genotypes within each of the three synthetic communities. Shaded areas indicate 95% confidence intervals of the correlation. Color codes in the bottom left box, on the right. **C.** Mean slope difference of the six *A. thaliana* genotypes used in this study. The slope difference indicates the effect of the genotype on the correlation between weight and isolate load - i.e. genotype * log_10_(cumulative isolate load) - following the model weight ∼ genotype * log_10_(cumulative isolate load) + experiment + error. Each treatment was analyzed individually, thus the model was utilized for each treatment separately. Kus3-1 was randomly selected as a reference. Dots indicate the medians, and vertical lines 95% credible intervals of the fitted parameter. Related to panel B. n=170 for PathoCom, n=151 for CommenCom, and n=182 for MixedCom. n=77-94 for the six *A. thaliana* genotypes.

**Figure S9.**
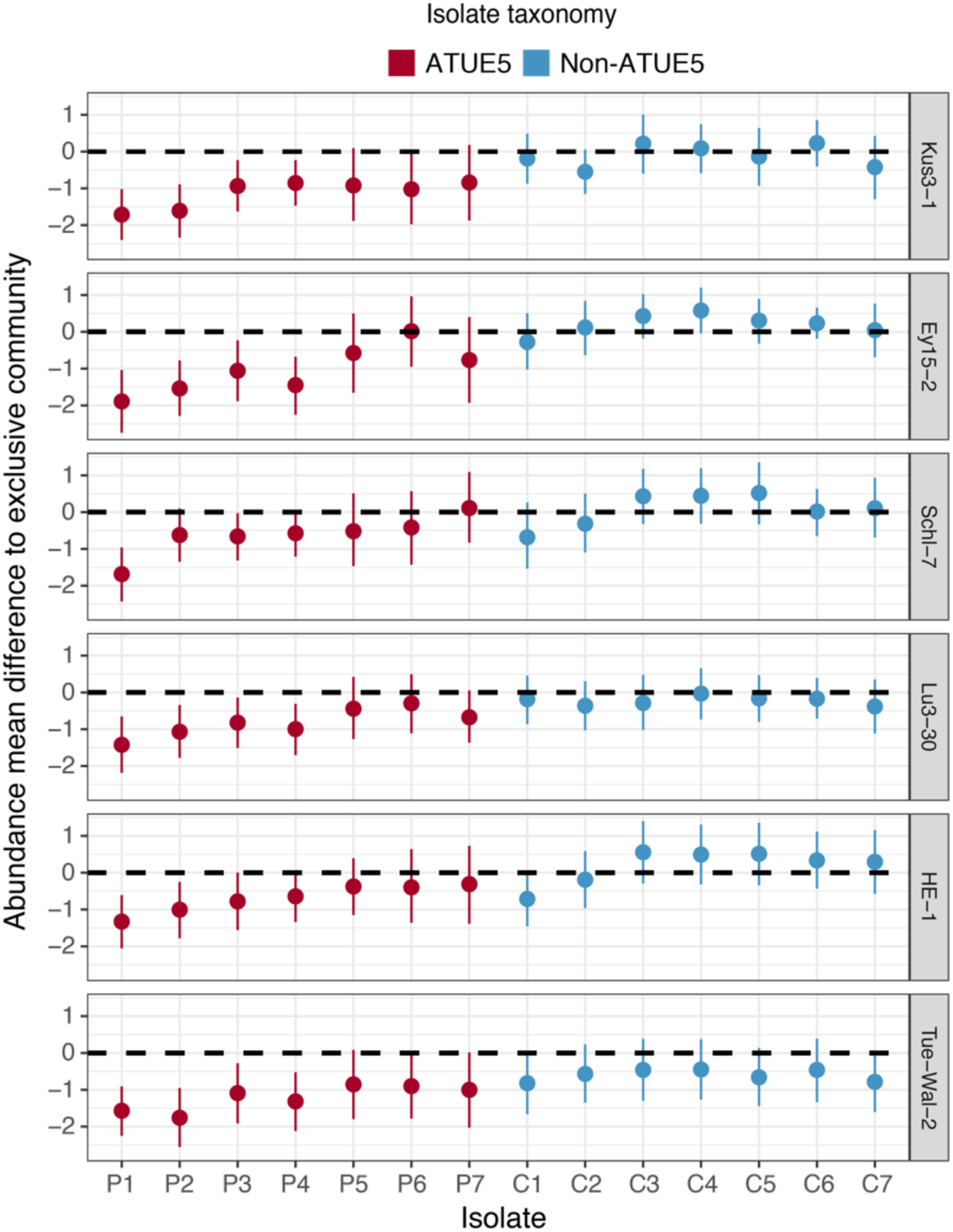
Effect of host genotype on abundance changes of the 14 barcoded isolates in MixedCom, when compared to their exclusive community (i.e., PathoCom for ATUE5 and CommenCom for non-ATUE5). Abundance effect mean differences were estimated with the model log_10_(isolate load) ∼ genotype * treatment * experiment + error for each individual strain. Thus, the genotype * treatment coefficient was estimated per each barcoded isolate. Dots indicate medians, and vertical lines 95% credible intervals of the fitted parameter.

**Figure S10.**
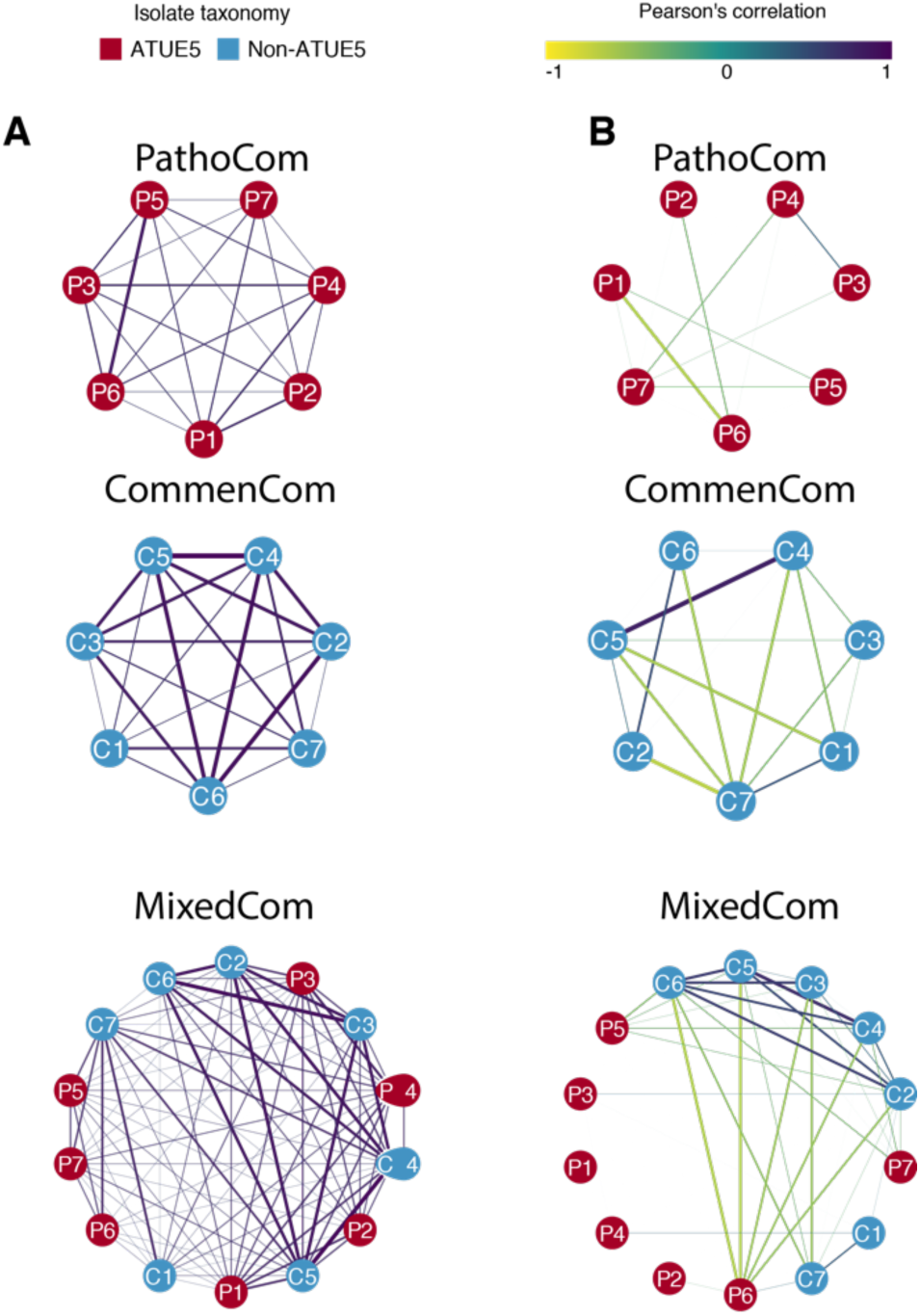
Correlation networks of barcoded bacteria. **A.** Correlation networks of absolute abundance in PathoCom, CommenCom and MixedCom. **B.** Correlation networks of relative abundance in PathoCom and CommenCom. Strengths of negative and positive correlations are indicated from yellow to purple. Boldness of lines is related to the strength of correlation, and only correlations > |±0.2| are shown. Node colors indicate the isolate classification: ATUE5 or non-ATUE5.

**Figure S11.**
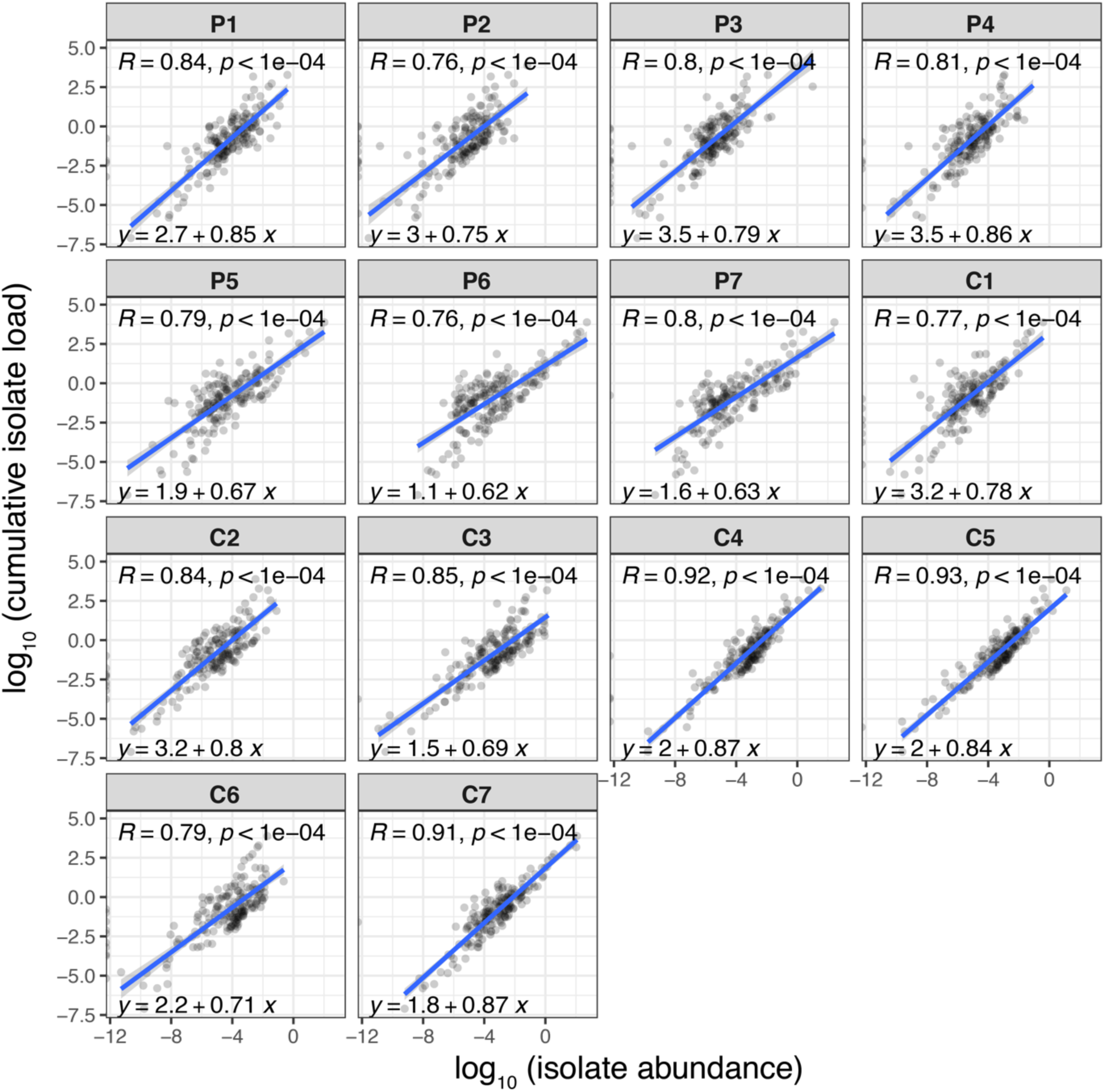
Correlations between the absolute abundance of each isolate and the cumulative bacterial abundance in MixedCom. Each panel represents an individual isolate. Pearson correlation (R) and p-value (p) are stated at the top, and the matching linear equation at the bottom of each panel. Shaded areas indicate 95% confidence intervals of the correlation curve.

**Figure S12.**
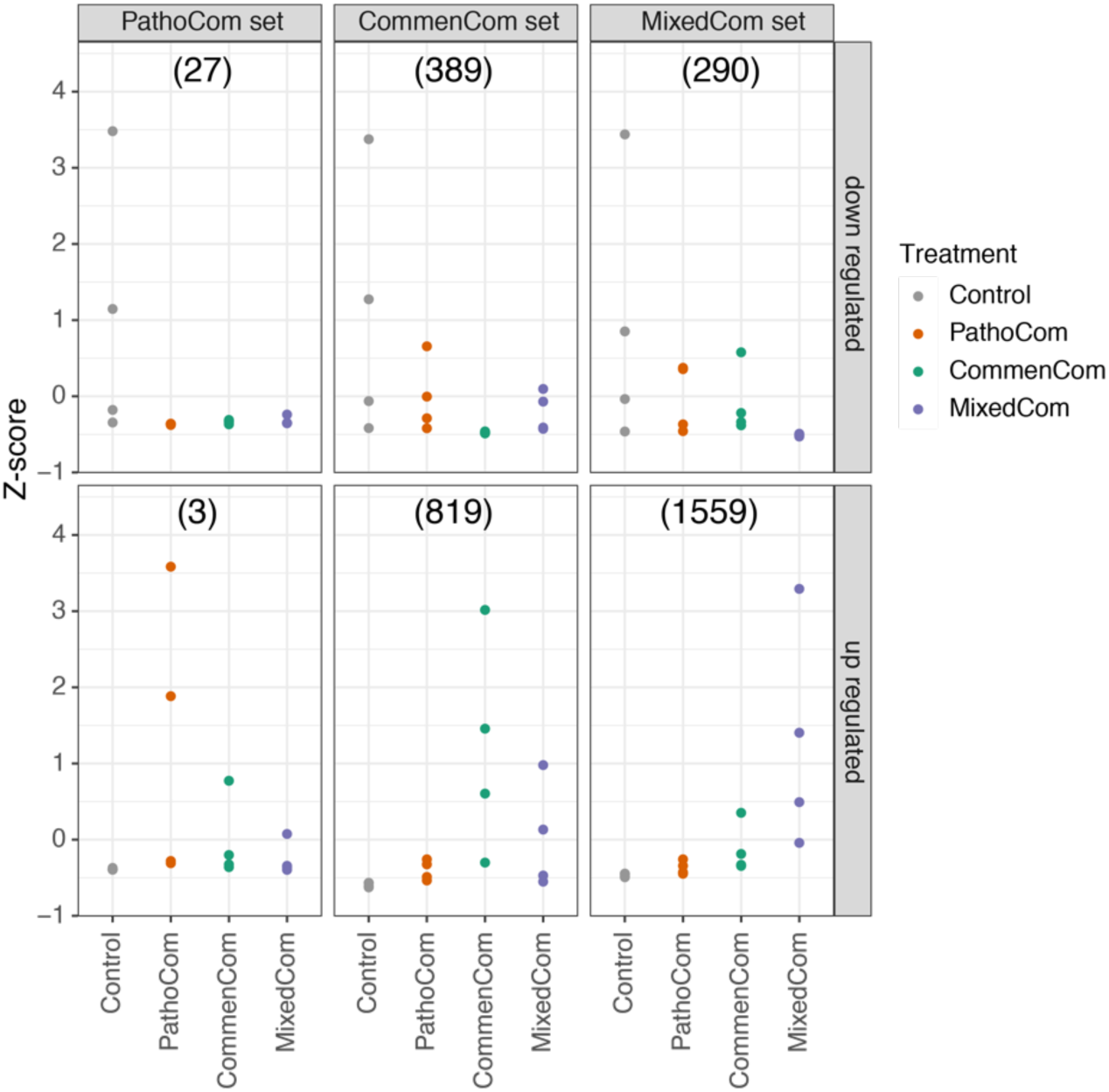
Comparison of PathoCom, CommonCom and MixedCom DEGs across treatments. The average z-score is presented for each sample. Downregulated and upregulated DEGs were analyzed separately. In brackets - the number of DEGs in each category. n=4.

**Figure S13.**
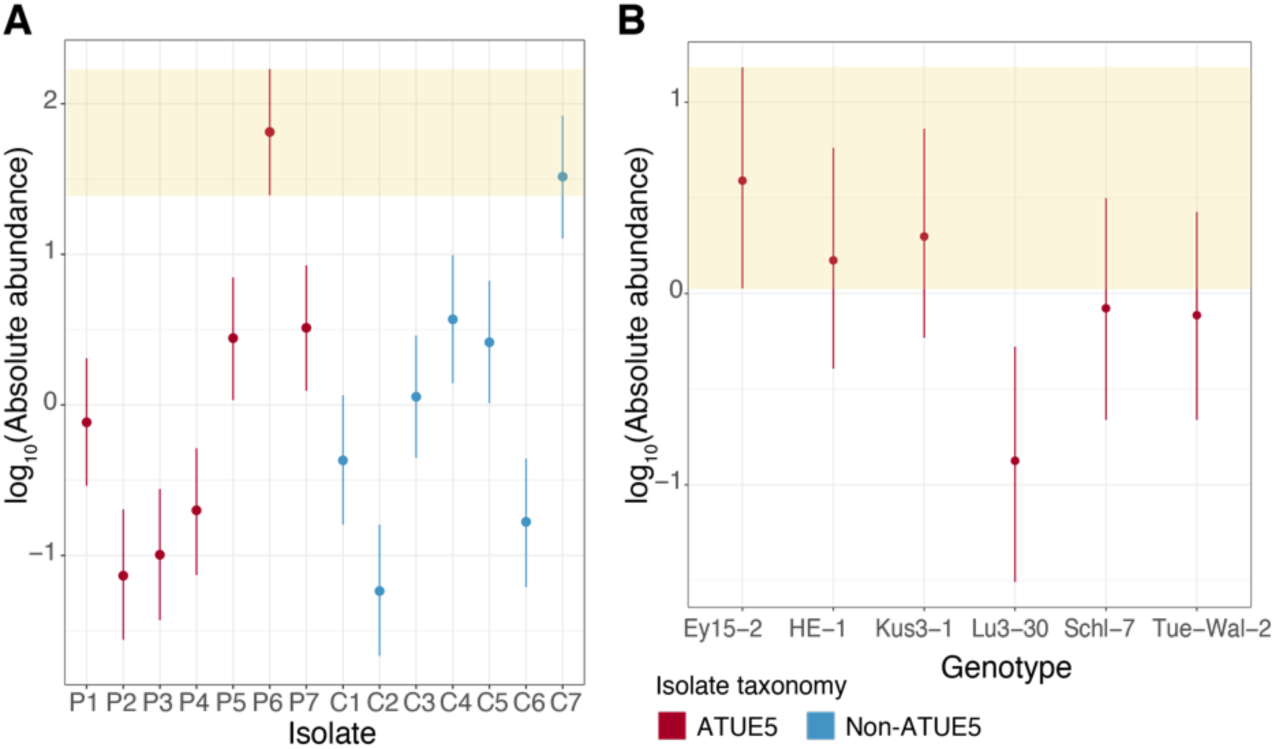
The abundance of P6 in MixedCom-infected hosts. **A.** Abundance of P6 compared with the other 13 barcoded bacteria in MixedCom-infected hosts, for all host genotypes. Dots indicate the medians, and vertical lines 95% credible intervals of the fitted parameter, following the model log_10_(isolate load) ∼ isolate * experiment + error. Shaded area denotes the 95% credible intervals of the isolate P6. **B.** The abundance of P6 in MixedCom-infected hosts, compared between the six *A. thaliana* genotypes used. Dots indicate the medians, and vertical lines 95% credible intervals of the fitted parameter, following the model log_10_(isolate load) ∼ genotype * experiment + error. Shaded area denotes the 95% credible intervals of the host genotype Ey15-2.

**Figure S14.**
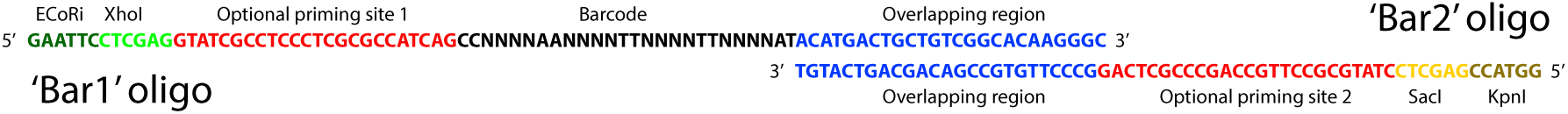
Illustration of barcodes design. Two single-stranded oligos were synthesized: ‘Bar1’ and ‘Bar2’. N symbolizes random nucleotides.

## Notes

### Competing Interest Statement

The authors have declared no competing interest.

### Summary of Updates

Updated version after correction of technical aspects (spacing and italics) + author name correction.

